# Whole-genome screening for near-diagnostic genetic markers for white oak species identification in Europe

**DOI:** 10.1101/2023.11.29.568959

**Authors:** Antoine Kremer, Adline Delcamp, Isabelle Lesur, Stefanie Wagner, Rellstab Christian, Erwan Guichoux, Thibault Leroy

## Abstract

**Context:** Identifying species in the European white oak complex has been a long standing concern in taxonomy, evolution, forest research and management. *Quercus petraea, Q. robur, Q. pubescens* and *Q. pyrenaica* are part of this species complex in western temperate Europe and hybridize in mixed stands, challenging species identification.

**Aims:** Our aim was to identify diagnostic single nucleotide polymorphisms (SNPs) for each of the four species that are suitable for routine use and rapid diagnosis in research and applied forestry.

**Methods:** We first scanned existing whole-genome and target-capture data sets in a reduced number of samples (training set) to identify candidate diagnostic SNPs, ie genomic positions being characterized by a reference allele in one species and by the alternative allele in all other species. Allele frequencies of the candidates SNPs were then explored in a larger, range-wide sample of populations in each species (validation step).

**Results:** We found a subset of 38 SNPs (ten for *Q. petraea*, seven for *Q. pubescens*, nine for *Q. pyrenaica* and twelve for *Q. robur*) that showed near-diagnostic features across their species distribution ranges with *Q. pyrenaica* and *Q. pubescens* exhibiting the highest and lowest diagnosticity, respectively.

**Conclusions:** We provide a new, efficient and reliable molecular tool for the identification of the species *Q. petraea, Q. robur, Q. pubescens* and *Q. pyrenaica*, which can be used as a routine tool in forest research and management. This study highlights the resolution offered by whole-genome sequencing data to design diagnostic marker sets for taxonomic assignment, even for species complexes with relatively low differentiation.

## 1. Introduction

Identifying species in the European white oak complex has been a long-standing concern in evolutionary biology as well as in forest research and management. According to the latest taxonomic classification, there are about fifteen oak species in Europe, which form the subsection of the Roburoids within the *Quercus* section (white oak section) (Denk *et al* 2017; Hipp *et al*, 2020). Within the continent, however, species richness varies, with higher species diversity in the Mediterranean region and in Eastern Europe compared to other areas (Camus, 1938; Le Hardy de Beaulieu and Lamant, 2006). In western temperate Europe, four white oaks species occur north of the Pyrenees and Alps (*Q. petraea, Q. robur, Q. pubescens* and *Q. pyrenaica*). Co-occurrence of all four species in the same forest is rare. The few reported cases indicate extensive gene flow and admixture between all four species, leading to considerable morphological variations and uncertainties when it comes to taxonomic classification based on morphological characters (Lepais *et al*, 2013; Leroy *et al*, 2017; Viscosi *et al*, 2009). The co-occurrence of the three species *Q. petraea*, *Q. robur* and *Q. pubescens* is more common, especially in the southern parts of the temperate range, for which hybridisation and morphological variation is well documented (Dupouey and Badeau, 1993; Grandjean and Sigaud, 1987; Macejovsky *et al*, 2020; Rellstab *et al*, 2016). Finally, forests with co-occurrences of two species and interspecific admixture have also raised questions about species classification. This is especially true for co-occurrences of *Q. petraea-Q. robur* (Bacilieri *et al*, 1995; Jurksiene and Baliuckas, 2014; Kelleher *et al*, 2005; Kremer *et al*, 2002; Yucedag and Gailing, 2013), but also for *Q. petraea-Q. pyrenaica* (Lopez de Heredia *et al*, 2009) and *Q. petraea-Q. pubescens* (Bruschi *et al*, 2000, Reutimann *et al,* 2020, 2023). This brief overview of species admixture and problems of taxonomic classification based on morphological characteristics highlights the pressing need for a time and cost efficient molecular tool for reliable species assignment within European white oaks for use in forest science and management.

In response to this challenge, molecular tools have been continuously improved and a number of species marker kits have been developed and applied during the last decade (Guichoux *et al,* 2011; Neophytou, 2014; Reutimann *et al*, 2020, Degen *et al*, 2021; Schroeder and Kersten, 2023). These methods have set new milestones for the delimitation of oak species, but their validity has been constrained by some biological and technical limitations. From a biological point of view, the markers used in the kits are still shared between the species, although interspecific differentiation of the selected markers was higher than in earlier studies. From a technical point of view, the genomic resources explored for selecting the marker candidates was very limited until recently. Using previously published genome-wide data and genome scans targeting genomic positions that maximise differentiation between populations of *Q. robur*, *Q. petraea*, *Q. pubescens* and *Q. pyrenaica*, we overcame these limitations and designed a new single-nucleotide (SNP) marker set for range-wide species identification in European white oaks.

Earlier genome scans for species differences showed that interspecific differentiation (*F_ST_*) followed an L-shaped distribution suggesting that there might be highly differentiated markers at an extremely low frequency within the genome (Reutimann *et al*, 2020; Scotti-Saintagne *et al*, 2004). Recent analysis of nucleotide diversity in genes underlying species barriers between European white oaks confirmed these expectations (Leroy *et al*, 2020b). Our approach built on these results by launching a systematic search of so-called species “diagnostic” SNPs within existing genome-wide resources. Ideally a diagnostic SNP contains a diagnostic allele of a given species that is fully fixed in that species and the alternate allele fixed in the other species. Earlier surveys (Scotti-Saintagne *et al*, 2004; Reutimann *et al*, 2020; Lesur *et al*, 2018) in European white oaks indicated that such ideal cases rarely exist. However, some markers exhibit species frequency profiles close to the ideal case (so-called near-diagnostic SNPs; for example an allele with a frequency larger than 0.9 in the target species, and alternate allele frequency larger than 0.9 in all other species) (Schroeder and Kersten, 2023). Only a few of such markers would then be enough to correctly assign trees to the correct species using appropriate analytical approaches. For example, Reutimann *et al* (2020) showed that five SNPs were enough for correctly classifying 95% of *Q. robur* reference trees, although the single SNPs were far from being diagnostic.

In this study we explored pool-sequenced whole-genome libraries of natural populations of four white oak species (*Q. petraea, Q. pubescens, Q. pyrenaica* and *Q. robur*) (Leroy *et al*, 2020b), and genome-wide capture-based sequences of *Q. petraea* and *Q. robur* (Lesur *et al*, 2018) to identify near-diagnostic SNPs for each of the four species. We describe the approaches and methods used to discover near-diagnostic SNPs, and explore the stability of diagnosticity over the distribution range of the four species.

Our main goal was to identify and validate a new set of near-diagnostic SNPs that can be used in the development of an efficient and cost-effective molecular tool for forest research and management. To this end, we focused on the variation of near-diagnostic SNPs across species, between populations within each species and between SNPs. We finally addressed the evolutionary drivers that may have contributed to the maintenance and/or modification of diagnosticity within the genome, and throughout the distribution range of the four species.

## 2. Material and Methods

### 2.1. Discovery of near-diagnostic markers

The discovery of near-diagnostic SNPs was conducted by scanning oak genomic data that have been generated in earlier studies assessing genomic diversity and differentiation in the four sympatric white oak species (*Quercus petraea, Q. pubescens, Q. pyrenaica, Q. robur;* Leroy *et al*, 2017 and 2020b, Lesur *et al*, 2018).

#### 2.1.1. Discovery of near-diagnostic SNPs in whole genome pool-sequenced (pool-seq) resources

##### 2.1.1.1. Pool sequencing

In Leroy *et al,* 2020b, we used leaf and bud samples from up to 20 adult trees of the four species coming from four different forests located at maximum 200 km away from each other in South West of France (Table 4 in Appendix). The sampled stands were of mixed oak composition (generally two or three species) and of natural origin. DNA extracts were pooled in equimolar amounts to obtain a single pool for each species. Libraries were then sequenced on nine to ten lanes for each of the four species (1 pool per species) on a Illumina HiSeq 2000 sequencing platform (Leroy *et al*, 2020b for details). In this study, to reduce the computation load, we only used two lanes per pool from SRA, namely ERR2215923 and ERR2215924, ERR2215937 and ERR2215938, ERR2215909 and ERR2215910, and ERR2215916 and

ERR2215917 for *Q. pubescens, Q. petraea*, *Q. pyrenaica* and *Q. robur* respectively. Raw reads were then trimmed using Trimmomatic (v. 0.33, Bolger *et al*, 2014) to remove low quality bases using the following parameters: LEADING:3 TRAILING:3 SLIDINGWINDOW:4:15 MINLEN:50.

##### 2.1.1.2. Mapping and SNP calling

Data from two sequencing lanes per species (from up to 10 lanes per species in Leroy *et al*, 2020b) were then mapped against the oak haplome assembly (“PM1N”, Plomion *et al*, 2018) using bwa mem (Li, 2013). PCR duplicates were removed using Picard v. 1.140 (http://broadinstitute.github.io/picard/). Samtools v.1.1 (Li, 2011) and PoPoolation2 v. 1.201 (Kofler *et al*, 2011) were then used to call bi-allelic SNPs with at least 10 reads of alternate alleles and a depth between 50 and 2000x at each position. To ensure a reasonably low rate of false positives due to Illumina sequencing errors, all SNPs with a MAF lower than 0.05 were discarded. A total of 24,345,915 SNPs were identified and then screened for their diagnostic value (see next paragraph).

##### 2.1.1.3. Genome scan for near-diagnostic SNPs

Allele frequencies were computed from the SNP-frequency-diff.pl script of PoPoolation2. SNPs exhibiting a high difference in allele frequency (Δ*p*>0.9 between the focal species and all other species) were then selected. All candidate diagnostic SNPs with a coverage lower than 80 in the four populations were discarded, in order to ensure that the high Δ*p* was not associated with inaccurate allele frequency estimation in low coverage regions. Despite the relatively limited linkage disequilibrium in oaks (Coq-Etchegaray *et al*, 2023) even in species barrier regions (Leroy *et al,* 2020b), the relatively high nucleotide diversity in oaks (Plomion *et al,* 2018; Saleh *et al*, 2022) allows several neighboring SNPs to be identified by this screening. We therefore selected the best SNPs per identified region considering the constraints associated with the SNP design (see below).

#### 2.1.2. Discovery of near-diagnostic SNPs in sequence-captured (seq-cap) genomic resources

In addition to the pool-seq resources, we mined a separate genome-wide resource that came from a sequence capture experiment of *Q. robur* and *Q. petraea* aiming at calling SNPs for inferring genomic relatedness among trees (Lesur *et al*, 2018). Here, the discovery population consisted of a far larger panel (245 adult trees in total) equally distributed between *Q. petraea* and *Q. robur* growing in the Petite Charnie forest located in the western part of France (Table 4 in Appendix). We used the capture data in complement of the pool-seq data to ensure a higher diagnosticity of the markers for this specific pair, given the larger panel of *Q. robur* and *Q. petraea* samples available in the capture data. The capture-based assay consisted in sequencing 2.9 Mb (15 623 target regions) on an Ion Proton System (Thermo Fisher, Scientific, Waltham, MA, USA) covering both genic and intergenic regions and resulted in the calling of more than 190,000 SNPs with a coverage of more than 10x (Lesur *et al*, 2018). The study provided allele frequencies of each SNP, and we screened the total set of SNPs for their differentiation between *Q. petraea* and *Q. robur*, by ranking their *F_ST_* values to complete the discovery panel. Although limited to two species (*Q. petraea* and *Q. robur*), this data set corresponded also to a genome-wide exploration of species differentiation implemented on a larger population sample (Lesur *et al*, 2018). It was therefore selected for this study, pending its relevance for selecting near-diagnostic markers for the remaining two species (*Q. pubescens* and *Q. pyrenaica*), which is investigated in this study.

### 2.2. Training and validation of near-diagnostic SNPs

#### 2.2.1. Training populations

The candidate diagnostic SNPs of the discovery panel were first tested on a limited number of oak individuals, with one sample per population for up to nine populations per species (19 to 48 samples per species) originating mainly from the western part of Europe (Table 5 in Appendix). The training experiment was conducted over two sessions that took place during two periods (training 1 and training 2, Table 5 in Appendix) with different samples (but from the same geographic range). The two sessions differed only by the samples included which was constrained by the availability of the material. The objective of the training step was to check whether the candidate SNPs exhibited near-diagnostic frequency profiles in natural populations originating mainly from the area of the discovery panel. The training step also included quality controls and repeatability assessments of the genotyping assay (see results paragraph 3.3.1).

#### 2.2.2. Validation populations

Given that the discovery and training of diagnostic SNPs was implemented on limited number of trees originating mostly from the western part of the distribution of the four species, we included a round of validation by increasing the sample sizes of the training populations and enlarging the collection of populations, studying the SNP diagnosticity across a larger part of the species’ natural distribution (Figure 1). Additionally, the validation step aimed also at reducing the number of SNPs, while still maintaining overall multilocus diagnosticity, in order to produce a low cost and easy to use screening tool in operational forestry. In total, 24 populations of *Q. petraea*, 10 of *Q. pubescens*, 6 of *Q. pyrenaica* and 19 of *Q. robur* were part of the validation set, representing in total 1,123 trees (Figure 1 and Table 5 in Appendix). All samples were collected in natural populations and their taxonomic status was assessed by the local collectors based on leaf morphology. Sampled populations were in most cases of mixed oak composition. Some of the populations were used in earlier large-scale genetic surveys (Gerber *et al*, 2014; Kremer *et al*, 2002), others were purposely collected for this study.

**Figure 1.**
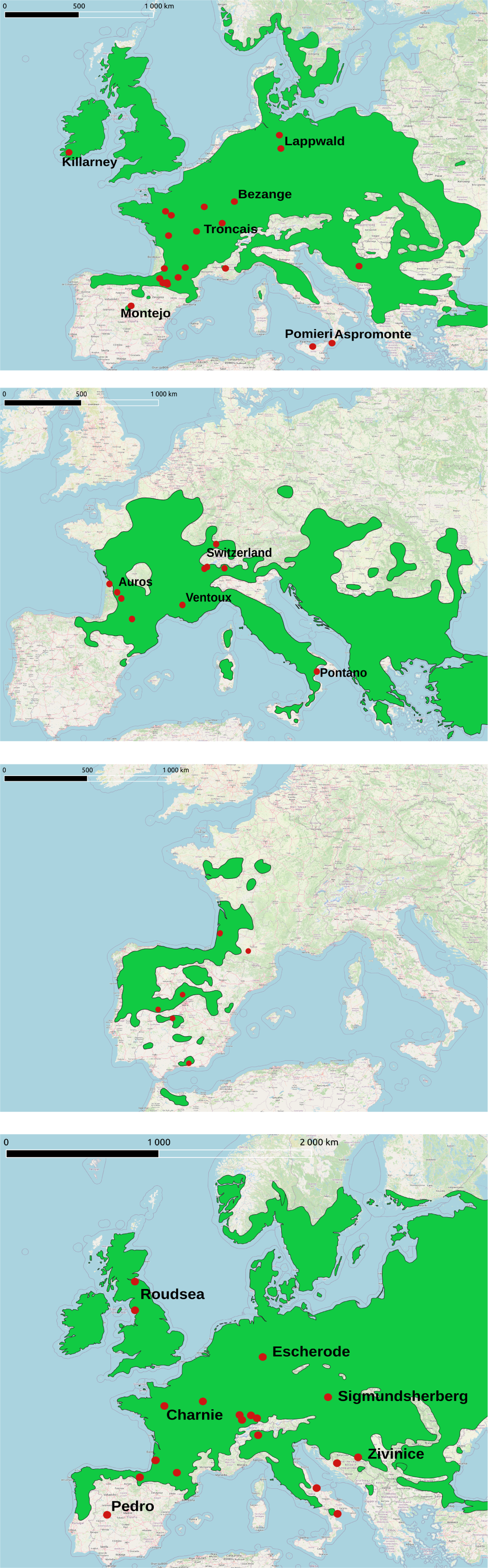
Geographic distribution of the validation populations. The green area corresponds to the distribution of the species according to Caudullo, Welk, et al. (2017). Red dots correspond to the origins of the validation populations. Populations identified by their name refer to populations for which frequency profiles of near-diagnostic alleles are later illustrated and discussed (paragaph 3.3.3)..

#### 2.2.3. Genotyping assay

Medium-throughput SNP genotyping assays were implemented on single tree DNA extracts using the MassARRAY® technology (Agena Bioscience, San Diego, CA, USA). The assay design, using the MassARRAY Assay Designer version 4.0.0.2, was performed on candidate SNPs from pool-seq and seq-cap resources. Nine multiplexes, for a total of 359 SNP (eight 40-plex and one 39-plex) were designed for identyfing the best markers. Genotyping was performed using iPLEX Gold chemistry following Ellis and Ong (2017) on a MassARRAY System CPM384 (Agena Biosciences) at the PGTB platform (doi:10.15454/1.5572396583599417E12). Data analysis was achieved using MassARRAY Typer Analyzer 4.0.4 (Agena Biosciences). After genotyping, we excluded all markers for which there was evidence that the candidate SNP identified during the discovery step was not recovered, for example when the SNP exhibited fixation across the four species at the same allele. We also discarded loci with weak (magnitude <5) or ambiguous signal (i.e. displaying more clusters than expected or unclear cluster delineation) and loci with more than 20% missing data. Following this selection process, 61 SNPs (in two multiplexes) were selected on the basis of their diagnostic value and their compatibility in one multiplex kit for subsequent genotyping on all the samples.

#### 2.2.4. Diagnosticity of candidate SNPs

Standard genetic statistics (allele frequencies, diversity statistics, differentiation and fixation indices) were estimated using GENEPOP (Raymond and Rousset, 1995) and ADEGENET software (Jombart, 2008).

We defined a metric of species diagnostic accuracy, which we coined « diagnosticity » index (*D*) to screen SNP alleles for their ability to be close to full diagnosticity.

Full diagnosticity requires two properties: fixation of the diagnostic allele in the target species and fixation of the alternate allele in the remaining species. These two properties are included in the metric *D.* Considering a set of *n* species, diagnosticity of an allele for species *x* (*D_x_*) regarding the remaining *(n-1)* species could be expressed as:

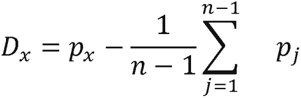

Where *p_x_* is the frequency of the candidate diagnostic allele in the target species *x,* and *p_j_* the frequency of the same allele in the alternate species *j*. *D_x_* amounts to the difference of allelic frequencies between species *x* and the remaining (*n-1*) species. *D_x_* is equivalent to the mean Gregorius genetic distance between species *x* and the three other species for a diallelic locus (Gregorius, 1984).

*D_x_*has two components, which account for the two properties of diagnosticity

- *p_x_*: the higher *p_x_*, the closer the near-diagnostic allele to fixation in the target species
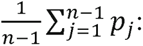 the lower the mean value of *p_j_*, the closer the alternate allele to fixation in the remaining (*n-1*) species.

*D_x_* is more appropriate for practical diagnostic assessments than the traditional differentiation metric *F_ST_* when more than two species are involved (see (Gregorius and Roberts, 1986) for a comparison of *D* and *F_ST_*). To illustrate the discrepancy between *D* and *F_ST_* regarding diagnosticity, consider the case of four species with frequency profiles (*p_1_*=1, *p_2_*=1, *p_3_*=0, *p_4_*=0). Addressing diagnosticity for species 1, *F_ST_* would yield 1, while *D_1_* would yield 0.67. *D_1_* accounts for the the lack of frequeny differences between species 1 and 2, while *F_ST_* does not.

By extension of the definition of a diagnostic allele, a near-diagnostic SNP is a SNP bearing near-diagnostic alleles, and diagnosticity of a species (or a population of that species) refers to the mean value of all near-diagnostic SNPs assessed for that species or population. Diagnosticity of candidate SNPs are estimated in the training and validation populations.

#### 2.2.5. Multilocus species clustering

To validate the selected near-diagnostic SNP for a multilocus species assignment procedure, we implemented an empirical clustering approach using Principal Component Analysis, free of any underlying evolutionary assumptions (ADEGENET, Jombart, 2008)). This method allows to check for the ability of the near-diagnostic SNPs to visually discriminate the 4 species.

## 3. Results

### 3.1. Discovery of near-diagnostic SNPs

All together we recovered 61 candidate near-diagnostic alleles, 49 originating from the pool-seq study, and 12 from the seq-cap analysis (Table 6 in Appendix). The candidate SNPs are distributed over all chromosomes (except chromosome 4) and their number ranges from 1 (chromosome 3, 9 and 12) to 17 (chromosome 2, Figure 2). In a few cases near-diagnostic markers of a given species clustered in pairs in a few spots (mainly for *Q. robur* on chromosome 2, 5, 6). In such cases one marker of the pair was discarded during the validation step. Near-diagnostic markers are distributed over 6 chromosomes for *Q. petraea, Q. pubescens* and *Q. pyrenaica*, and over 8 chromosomes for *Q. robur*. As indicated by their location on the chromosomes, the minimum physical distance of near-diagnostic SNPs located on the same chromosomes was 17 Kb (Table 6 in Appendix). All except two SNPs are located on scaffolds that are anchored on the pseudo-chromosome assembly of the oak genome as shown in Figure 2.

**Figure 2.**
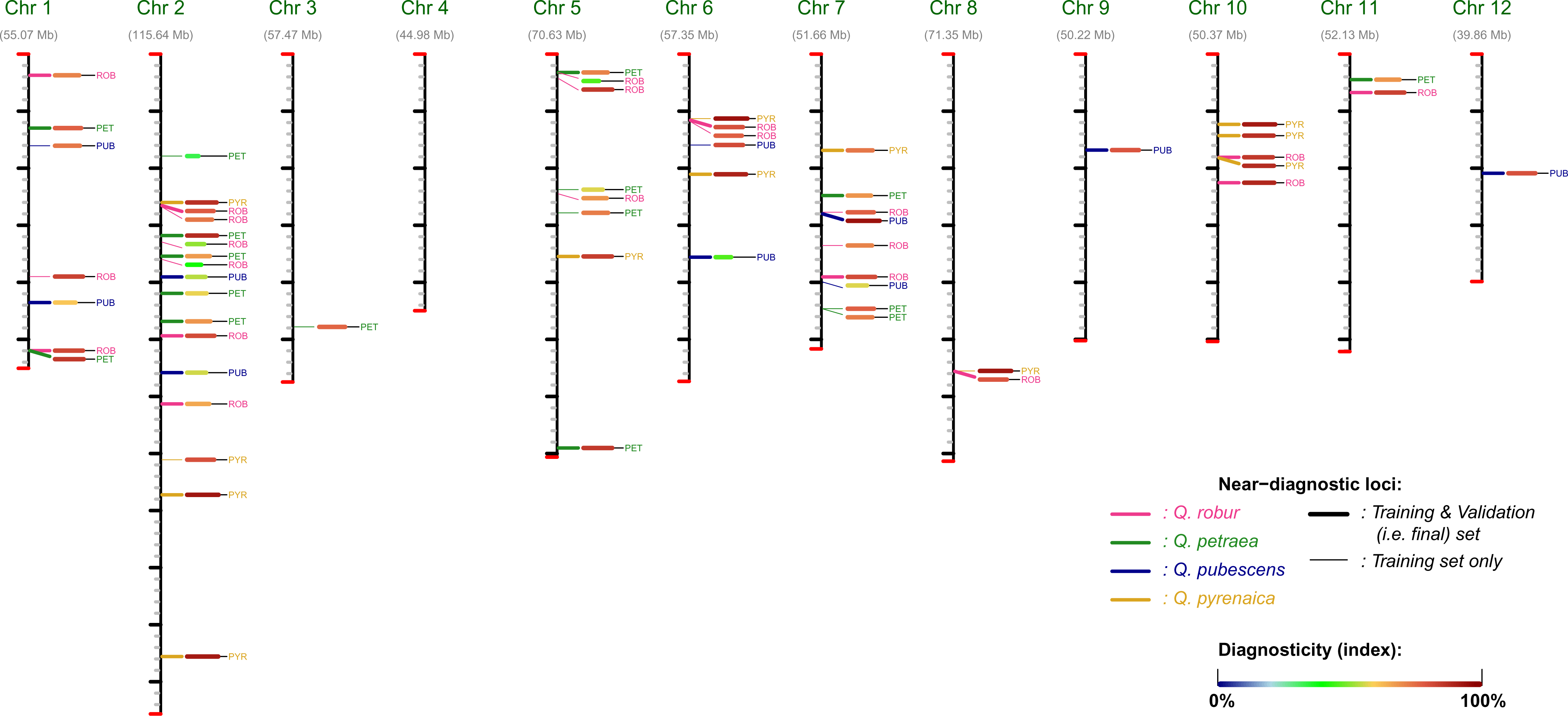
Genomic location of the near-diagnostic SNPs on the 12 oak (pseudo-)chromosomes of the oak chromosome. The color code of the marker corresponds to the species name for which the SNP is expected to be diagnostic (our design), with *Q. robur, Q. petraea, Q. pubescens and Q. pyrenaica*, shown in pink, green, blue and yellow, respectively. For each SNP, the diagnosticity of each marker at the training stage is indicated following the proportional and color scale shown. Thin and bold lines both indicate the location of the SNPs, but separates SNPs that were excluded or included in our final set of 38 SNPs, respectively. Note that two diagnostic SNPs are not shown since they are located on scaffolds that are not anchored on the oak pseudochromosomes (see Table 6 in Appendix).

### 3.2. Diagnosticity of candidate SNPs in the training set

The 61 candidate near-diagnostic SNPs exhibited allele frequency profiles close to the requisite properties of a diagnostic SNP but did not fulfill entirely criteria of full diagnosticity (Figure 2, Figure 6 in Appendix). *D* values indeed varied between 0.283 and 0.963. Most of the near-diagnostic SNPs (92%, 56/61) exhibit *D* scores greater than 0.50 (mean value 0.758). Among the 61 SNPs, 16 are candidate diagnostic of *Q. petraea*, 11 of *Q. pubescens*, 12 of *Q. pyrenaica* and 22 of *Q. robur*.

Diagnosticity scores were higher in the pool-seq uncovered set (*D*=0.771) than in the seq-cap uncovered set (*D*=0.704).

Concerning the near-diagnostic SNPs identified with the pool-seq data, diagnosticity was highest for *Q. pyrenaica* (0.897) and *Q. robur* (0.780) and lower in *Q. petraea* (0.736) and *Q. pubescens* (0.657). Deviations to full diagnosticity in the two latter species are associated with different patterns (Figure 6 in Appendix).:

● Lower diagnosticity in *Q. petraea* was mostly related to the sharing of the diagnostic allele with the other species, especially with *Q. pubescens*.
● Lower diagnosticity for *Q. pubescens* was mainly due to three SNPs (Sc0000170_630013, Sc0000192_329301 and Sc0000482_334917) that showed substantial deviation from fixation within *Q. pubescens* (frequency being respectively 0.468, 0.587, 0.283) while the alternate alleles were fixed in the three other species.

Concerning the seq-cap uncovered SNPs, we selected 12 SNPs that exhibited the highest species differentiation in the Petite Charnie population. As expected, all 12 SNPs showed strong frequency differences between *Q. petraea* and *Q. robur* in our training panel. Eight out of the 12 SNPs exhibited allele frequency differences among the four species consistent with diagnosticity requirements for four species, with the near-diagnostic marker being almost fixed in the reference diagnostic species and present at very low frequencies in all the three remaining species (Figure 6 in Appendix). The four remaining candidate SNPs exhibited near-diagnostic alleles being almost fixed, not only in one but in two species:

● Sc0000040_1694351 in *Q. petraea* and *Q. pubescens*
● Sc0000481_366275 in *Q. robur* and *Q. pyrenaica*
● Sc0000546_456229 in *Q. robur* and *Q. pyrenaica*
● Sc0000598_295142 in *Q. robur* and *Q. pyrenaica*

### 3.3. Validation of the near-diagnostic SNPs

#### 3.3.1. Screening of near-diagnostic SNPs

The validation step aimed at verifying the diagnosticity of the candidate SNPs on a larger geographic scale, while at the same time optimizing the assay by selecting the best SNPs according to various genetic and technical criteria. We thus attempted to optimize the MassARRAY® genotyping assays by reducing the number of near-diagnostic SNPs and combine them in one final assay, without limiting the species assignment purpose and reducing its diagnosticity. Indeed given the frequency profiles of near-diagnostic alleles we observed in the training set (Figure 6 in Appendix), the required number of near-diagnostic SNPs for species assignment can be limited to a handful of markers (Reutimann *et al*, 2020). We aimed at selecting about 10 near-diagnostic SNPs per species for the final design of the operational assay. The following criteria were applied (Table 6 in Appendix):

● Repeatability and clarity of the cluster delimitation on the scatter plots
● Diagnosticity of SNPs
● A nearly equal numbers of near-diagnostic SNPs per species

Combining the remaining SNPs within one or two multiplex sets, resulted in amplification incompatibilities among SNPs which lead us to discard additional SNPs. Finally a total of ten near-diagnostic SNPs were selected for *Q. petraea*, seven for *Q. pubescens*, nine for *Q. pyrenaica* and twelve for *Q. robur* (Table 6 in Appendix).

#### 3.3.2. Allele frequency profiles of near-diagnostic SNPs in the validation populations

Overall, the average diagnosticity of the 38 near-diagnostic SNPs was slightly higher in the validation than in the training populations, with the exception of *Q. pyrenaica* (Figure 3, Figure 6 in Appendix): 0.784 (validation) *vs* 0.715 (training) in *Q. petraea,* 0.747 *vs* 0.690 in *Q. pubescens*, 0.876 *vs* 0.897 in *Q. pyrenaica,* 0.841 *vs* 0.758 in *Q. robur.* The lower diagnosticity of *Q. pyrenaica* in the validation set (*vs* the training set) was due to SNP Sc0000307_852597, which exhibited contrasting values between the training (0.753) and validation set (0.546) (Table 7 in Appendix).

**Figure 3.**
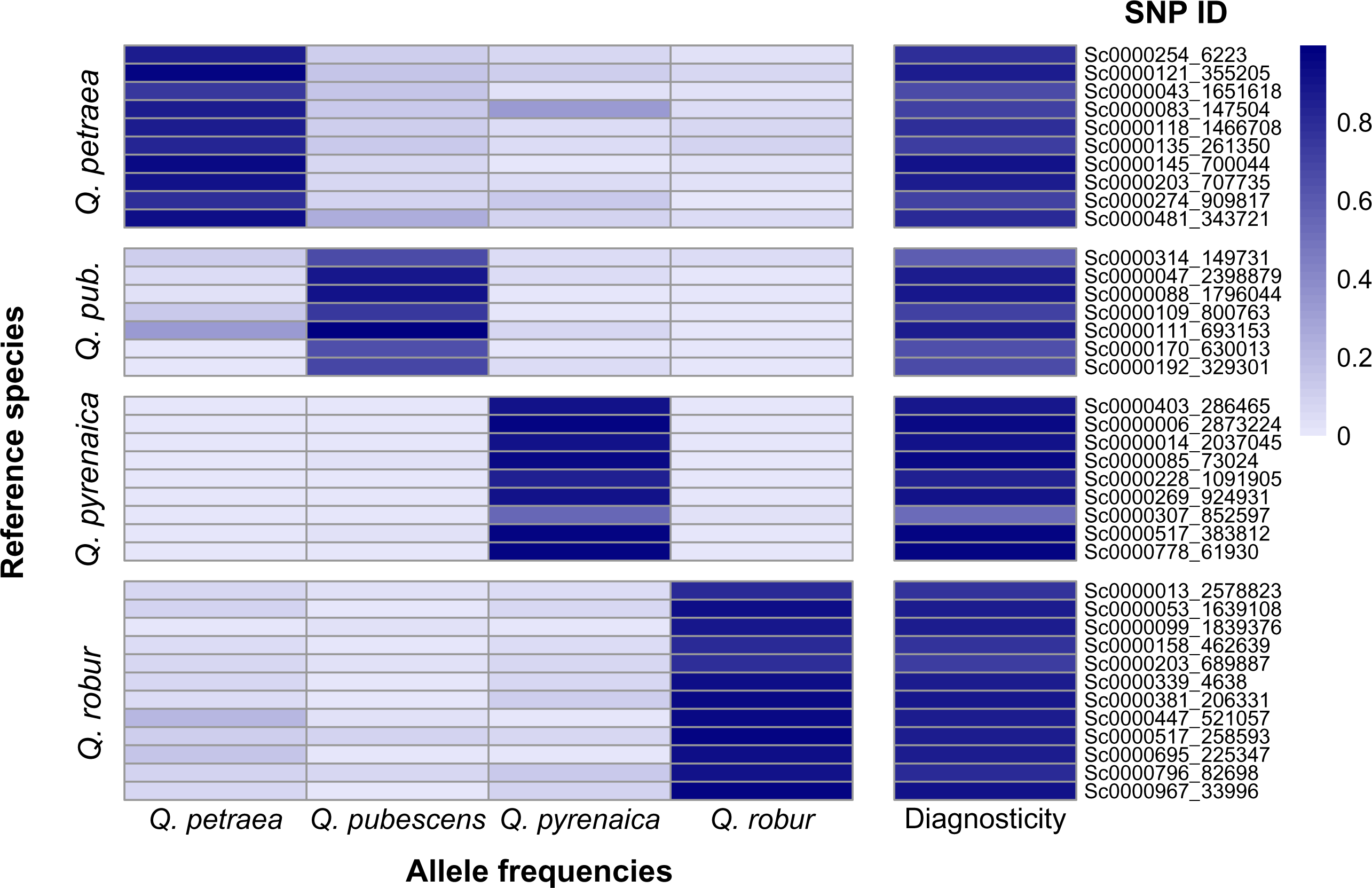
Heat map of frequencies of near-diagnostic alleles and diagnosticity in the validation populations of the four species. SNPs are clustered for their diagnostic value for each species (reference species): First to fourth columns correspond respectively to near-diagnostic alleles of *Q*. *petraea*, *Q.pubescens*, *Q. pyrenaica* and *Q. robur*.

However, the validation populations provided the opportunity to explore the stability of the allele frequency profiles across geographic regions, and thus addressed the maintenance of diagnosticity of individual SNPs across the distribution of the four species. Most near-diagnostic SNPs exhibited larger genetic differentiation between populations within a given species than usually found (Scotti-Saintagne *et al*, 2004) in oak species (Table 1, 2, 3). Mean intraspecific *F_ST_* values of near-diagnostic SNPs amounted to 0.104, 0.192, 0.042 and 0.104 for *Q. petraea, Q. pubescens, Q. pyrenaica,* and *Q. robur*, respectively. Furthermore, *F_ST_* values within a species exhibited large variation among SNPs. For example, *F_ST_* values of near-diagnostic SNPs of *Q. petraea* between *Q. petraea* populations varied between 0.012 and 0.252. *Quercus pyrenaica* is an exception to these general rules (0.042), as the mean *F_ST_* is much lower than for the 3 other species and the range of variation reduced (−0.022 to 0.142, data not shown).

**Table 1.**
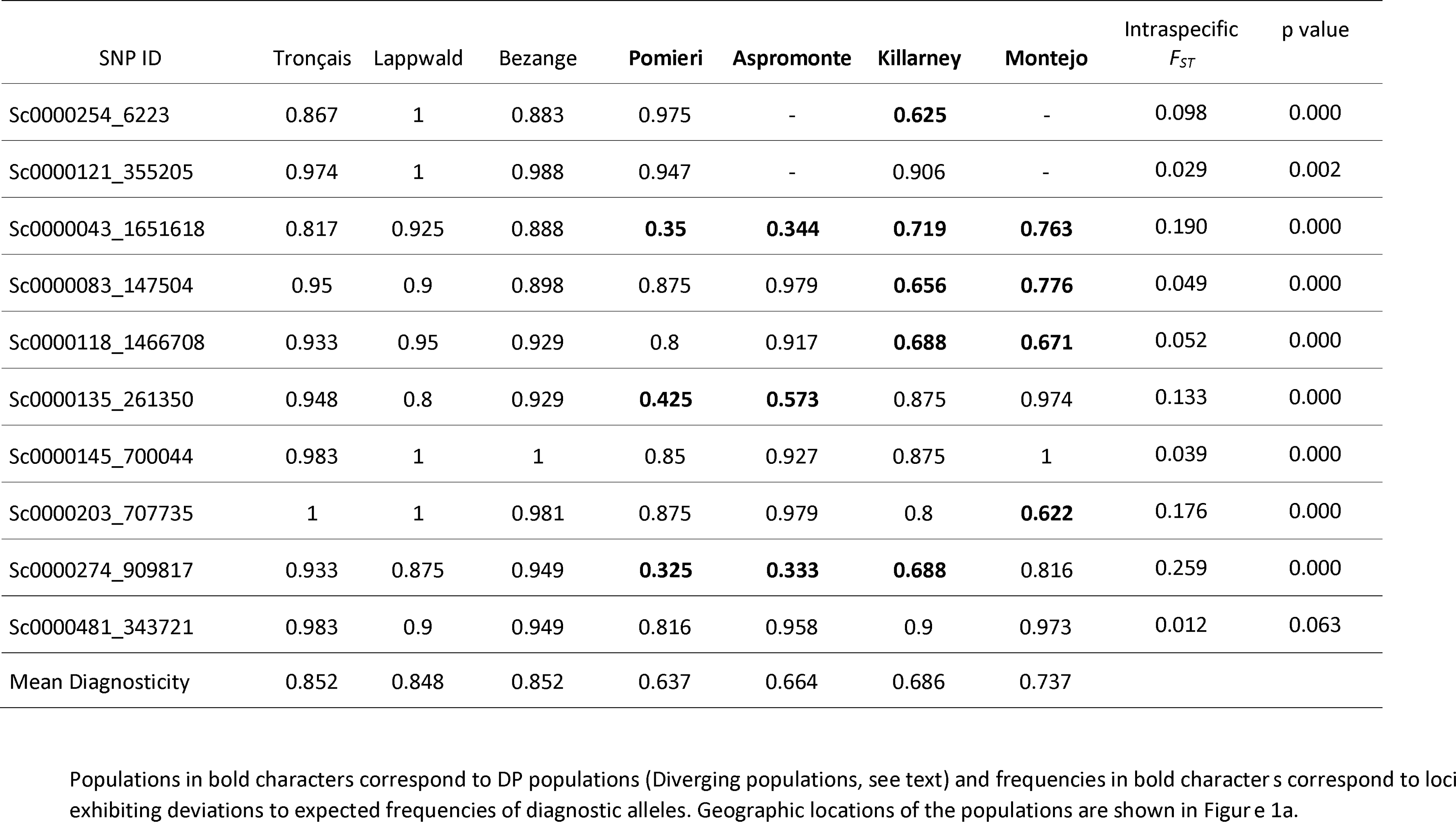
Frequencies and differentiation of near-diagnostic alleles of Q. petraea in Q. petraea populations.

**Table 2.** Frequencies and differentiation of near-diagnostic alleles of Q. pubescens in Q. pubescens populations.

| SNP ID | Auros | Pantano | Switzerland<sup>a</sup> | Ventoux | Intraspecific<br>$F_{ST}$ | P<br>value |
| --- | --- | --- | --- | --- | --- | --- |
| Sc0000314_149731 | 0.821 | - | **0.531** | **0.521** | 0.113 | 0.003 |
| Sc0000047_2398879 | 0.946 | 1.000 | 0.694 | 0.868 | 0.127 | 0.000 |
| Sc0000088_1796044 | 1.000 | 1.000 | 0.806 | 0.812 | 0.128 | 0.000 |
| Sc0000109_800763 | 0.839 | 0.882 | **0.528** | **0.692** | 0.096 | 0.000 |
| Sc0000111_693153 | 1.000 | 0.987 | 1.000 | 1.000 | -0.004 | 0.682 |
| Sc0000170_630013 | 0.907 | 0.987 | **0.222** | **0.400** | 0.506 | 0.000 |
| Sc0000192_329301 | 0.880 | 0.986 | **0.333** | **0.487** | 0.376 | 0.000 |
| Mean Diagnosticity | 0.865 | 0.926 | **0.540** | **0.635** |  |  |

<sup>a</sup>Switzerland population assembles data of populations Ayent, Cordola, Remigen and Saillon described in Table 5 in Appendix.

**Table 3 in Appendix.** Frequencies of near-diagnostic alleles and differentiation of Q. robur in Q. robur populations.

| SNP ID | Zivinice | Sigmunds-herberg | Charnie | Escherode | Pedro | Roudsea<sup>a</sup> | Intraspecific $F_{ST}$ | P value |
| --- | --- | --- | --- | --- | --- | --- | --- | --- |
| Sc0000013_2578823 | 0.711 | 0.925 | 0.875 | 0.750 | **0.273** | **0.553** | 0.140 | 0.000 |
| Sc0000053_1639108 | 1.000 | 1.000 | 1.000 | 0.857 | 0.818 | **0.500** | 0.098 | 0.009 |
| Sc0000099_1839376 | 0.868 | 0.868 | 0.889 | 0.972 | 0.818 | 0.711 | 0.040 | 0.000 |
| Sc0000158_462639 | 0.763 | 0.975 | 0.889 | 0.806 | 0.864 | **0.658** | 0.071 | 0.000 |
| Sc0000203_689887 | 0.842 | 0.750 | 1.000 | 0.806 | 0.955 | **0.632** | 0.095 | 0.000 |
| Sc0000339_4638 | 1.000 | 1.000 | 0.944 | 0.941 | **0.227** | 0.853 | 0.359 | 0.000 |
| Sc0000381_206331 | 1.000 | 1.000 | 1.000 | 1.000 | **0.500** | **0.737** | 0.233 | 0.000 |
| Sc0000447_521057 | 1.000 | 0.950 | 0.944 | 1.000 | 1.000 | 0.816 | 0.042 | 0.001 |
| Sc0000517_258593 | 0.938 | 0.975 | 0.938 | 0.917 | 1.000 | 0.974 | -0.003 | 0.166 |
| Sc0000695_225347 | 0.947 | 1.000 | 0.889 | 0.917 | **0.773** | **0.763** | 0.050 | 0.000 |
| Sc0000796_82698 | 0.842 | 0.925 | 0.944 | 0.972 | 0.864 | 0.789 | 0.021 | 0.016 |
| Sc0000967_33996 | 1.000 | 1.000 | 1.000 | 1.000 | 1.000 | **0.763** | 0.106 | 0.000 |
| Mean Diagnosticity | 0.851 | 0.889 | 0.885 | 0.854 | 0.700 | 0.671 |  |  |

<sup>a</sup>Roudsea population assembles data of populations Dalkeith and Roudsea described in Table 5 in Appendix.

**Table 4a in Appendix.**
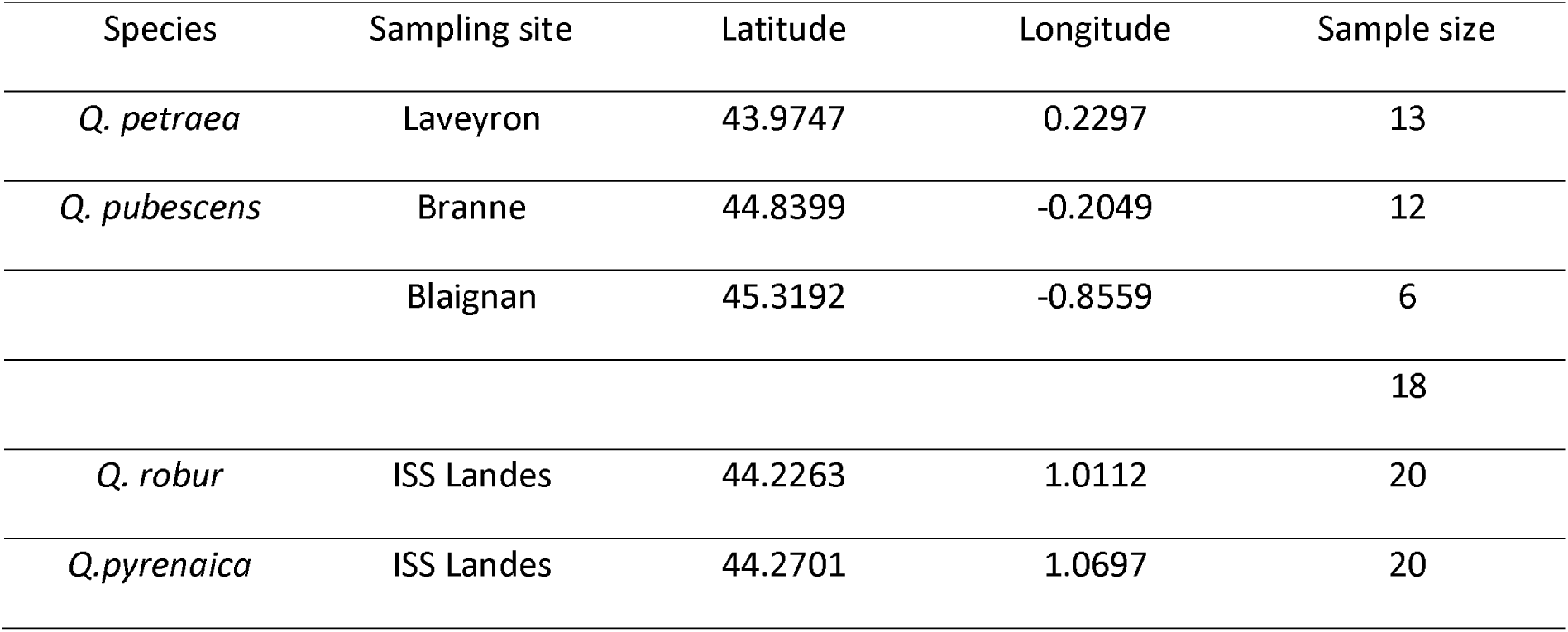
Discovery samples of the whole genome pool-sequenced resources.

**Table 4b in Appendix.** Discovery samples of the sequence captured genomic resources.

| Species | Sampling site | Latitude | Longitude | Sample Size |
| --- | --- | --- | --- | --- |
| *Q. petraea* | La Petite Charnie | 48.086 | -0.168 | 110 |
| *Q. robur* | La Petite Charnie | 48.086 | -0.168 | 135 |

**Table 5 in Appendix.**
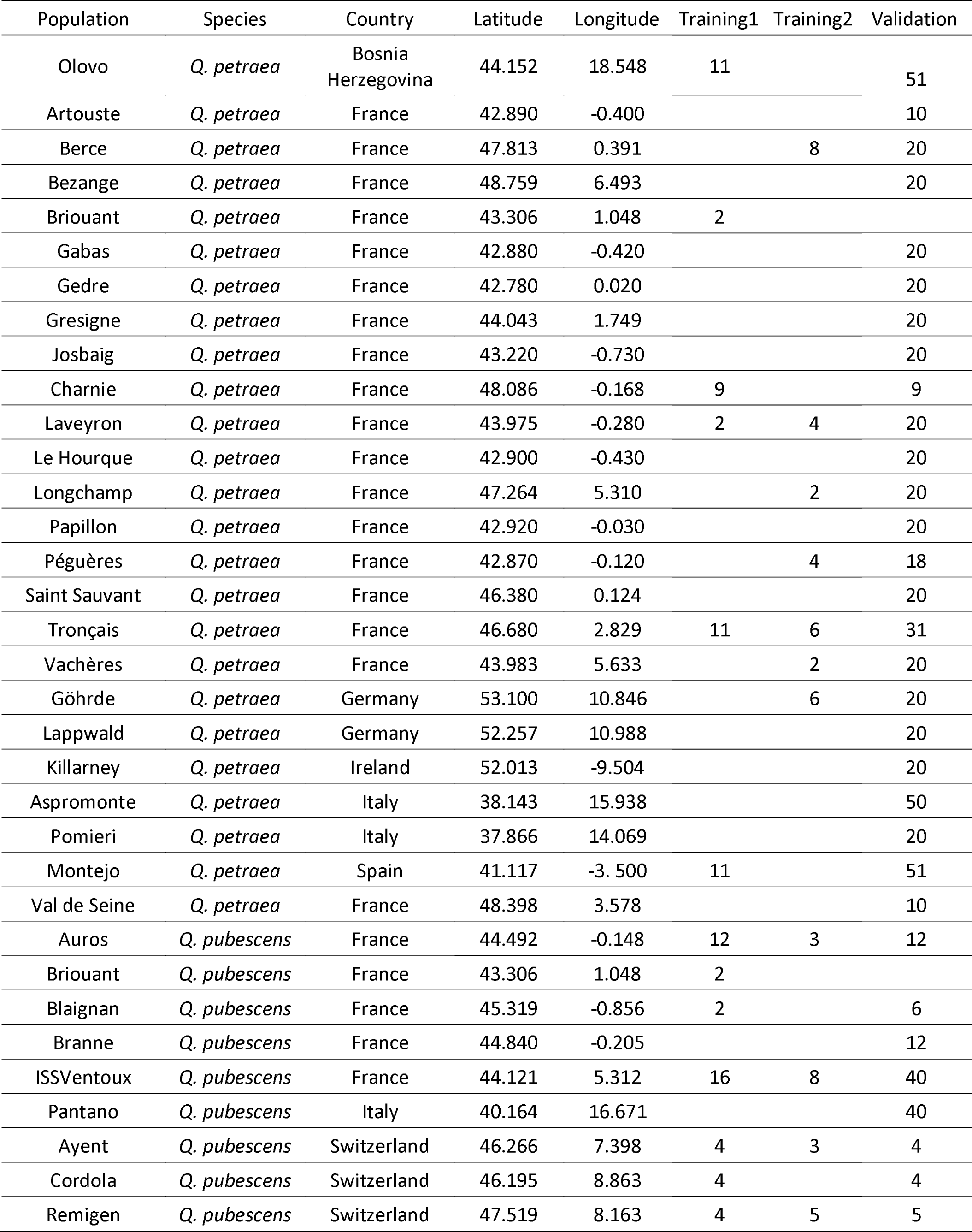
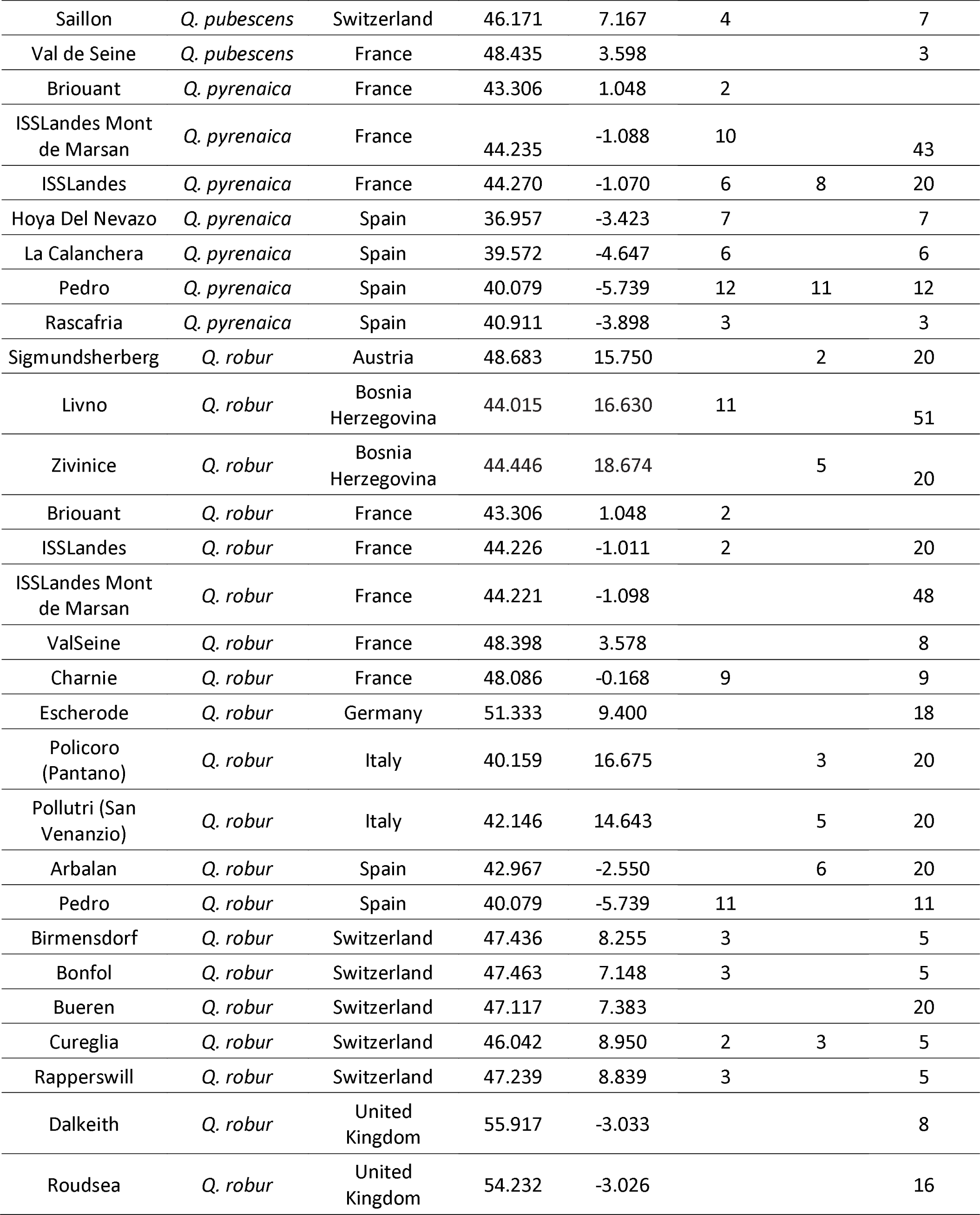
Geographic origins of training and validation samples.

**Table 6 in Appendix.**
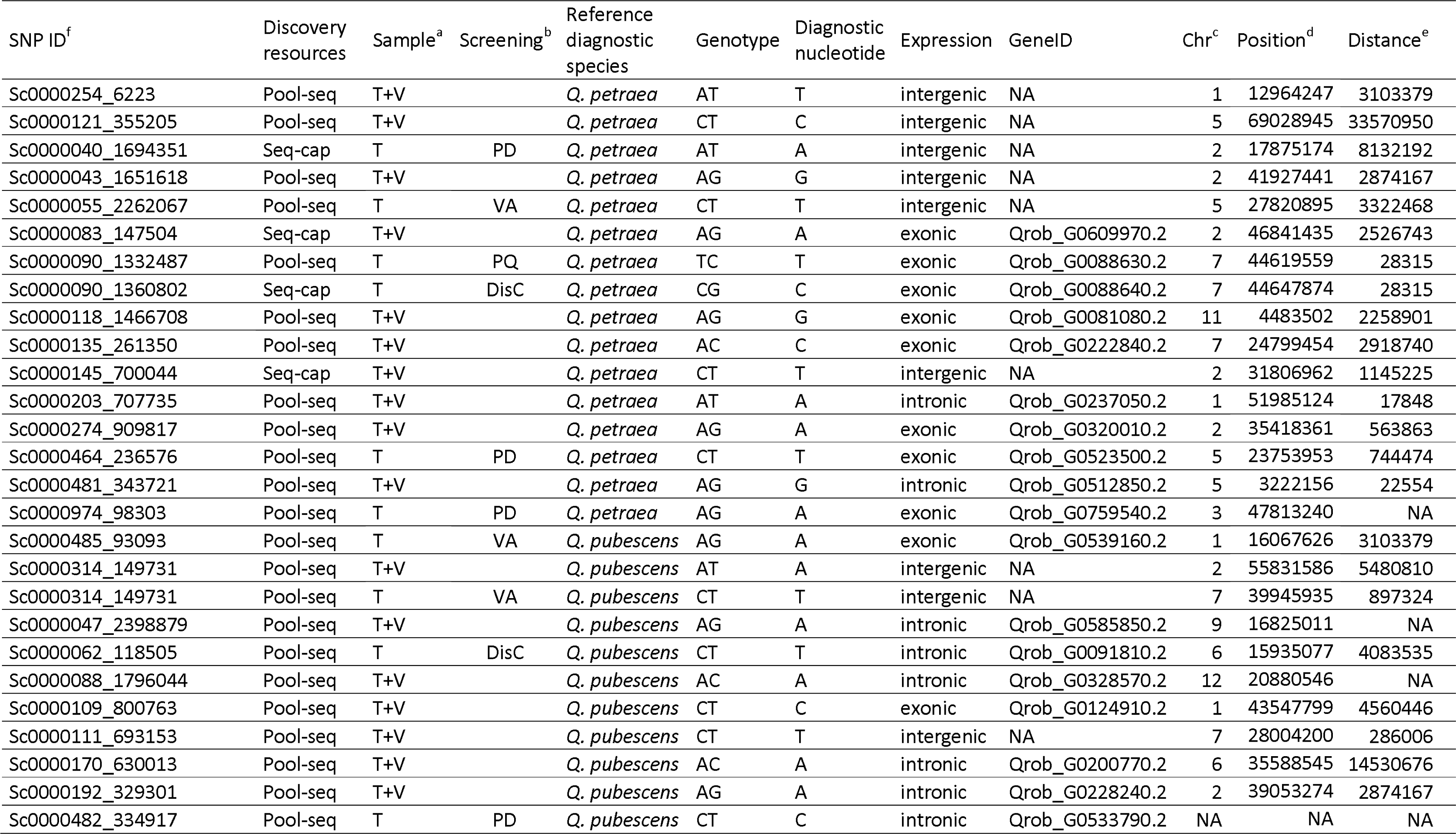
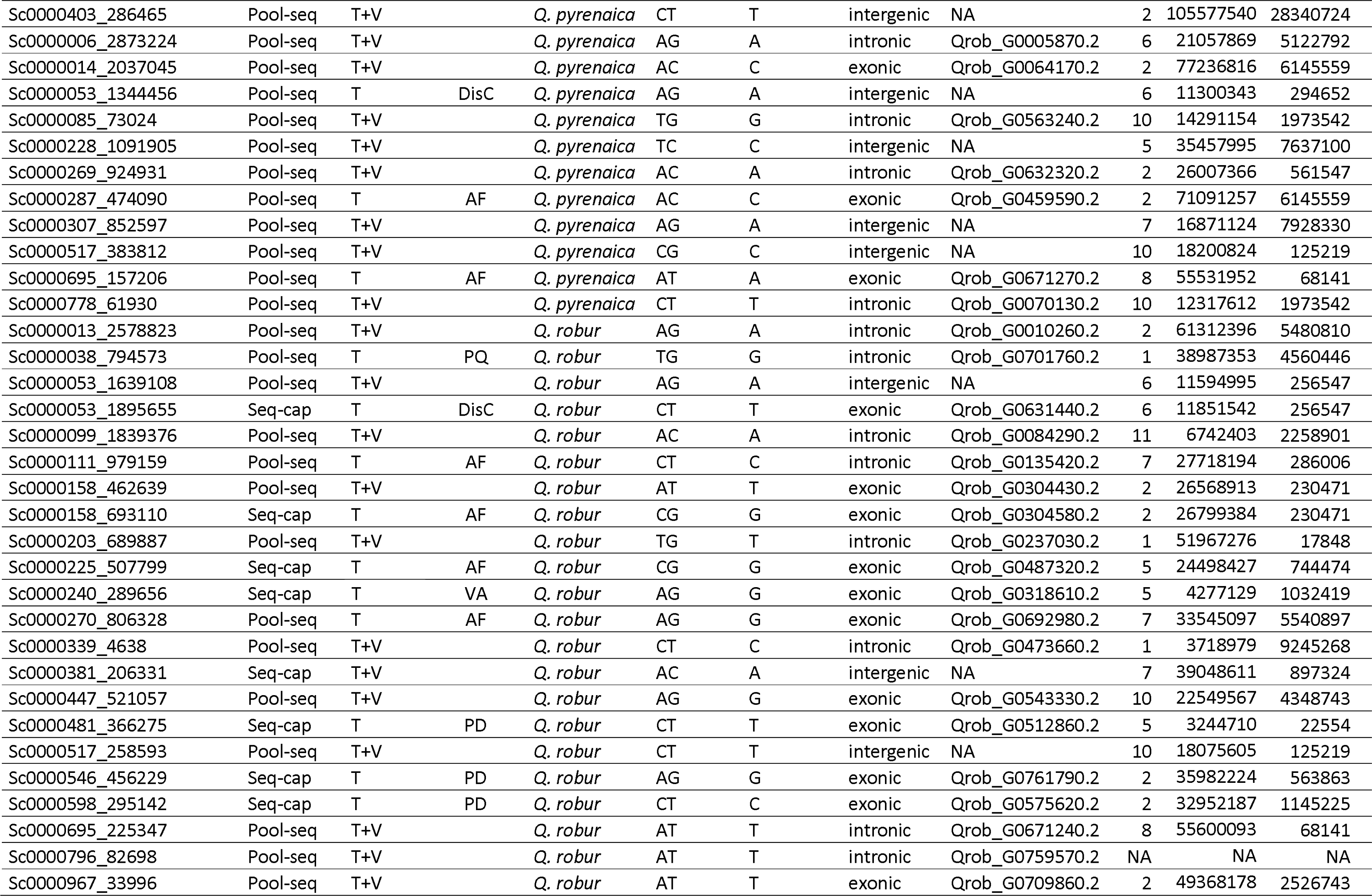
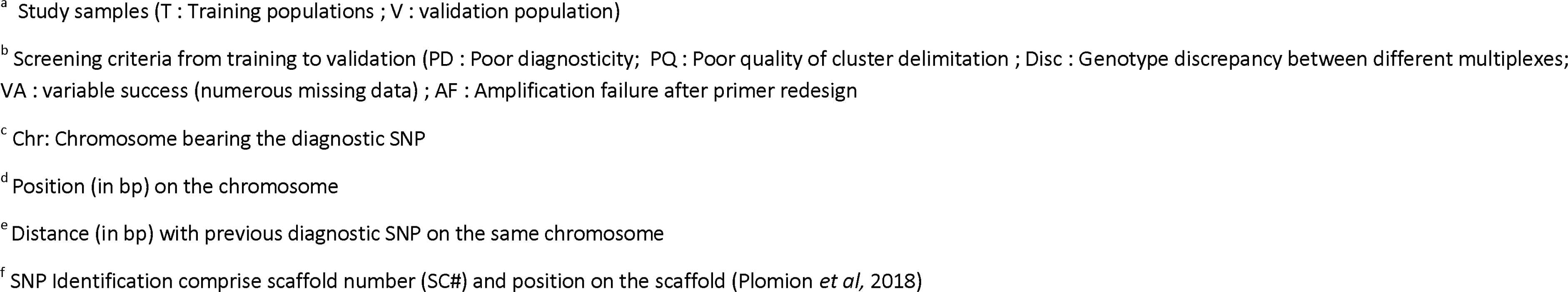
Genetic and genomic features of near-diagnostic SNPs.

| SNP ID<sup>f</sup> | Discovery resources | Sample<sup>a</sup> | Screening<sup>b</sup> | Reference diagnostic species | Genotype | Diagnostic nucleotide | Expression | GeneID | Chr<sup>c</sup> | Position<sup>d</sup> | Distance<sup>e</sup> |
| --- | --- | --- | --- | --- | --- | --- | --- | --- | --- | --- | --- |
| Sc0000254_6223 | Pool-seq | T+V |  | *Q. petraea* | AT | T | intergenic | NA | 1 | 12964247 | 3103379 |
| Sc0000121_355205 | Pool-seq | T+V |  | *Q. petraea* | CT | C | intergenic | NA | 5 | 69028945 | 33570950 |
| Sc0000040_1694351 | Seq-cap | T | PD | *Q. petraea* | AT | A | intergenic | NA | 2 | 17875174 | 8132192 |
| Sc0000043_1651618 | Pool-seq | T+V |  | *Q. petraea* | AG | G | intergenic | NA | 2 | 41927441 | 2874167 |
| Sc0000055_2262067 | Pool-seq | T | VA | *Q. petraea* | CT | T | intergenic | NA | 5 | 27820895 | 3322468 |
| Sc0000083_147504 | Seq-cap | T+V |  | *Q. petraea* | AG | A | exonic | Qrob_G0609970.2 | 2 | 46841435 | 2526743 |
| Sc0000090_1332487 | Pool-seq | T | PQ | *Q. petraea* | TC | T | exonic | Qrob_G0088630.2 | 7 | 44619559 | 28315 |
| Sc0000090_1360802 | Seq-cap | T | DisC | *Q. petraea* | CG | C | exonic | Qrob_G0088640.2 | 7 | 44647874 | 28315 |
| Sc0000118_1466708 | Pool-seq | T+V |  | *Q. petraea* | AG | G | exonic | Qrob_G0081080.2 | 11 | 4483502 | 2258901 |
| Sc0000135_261350 | Pool-seq | T+V |  | *Q. petraea* | AC | C | exonic | Qrob_G0222840.2 | 7 | 24799454 | 2918740 |
| Sc0000145_700044 | Seq-cap | T+V |  | *Q. petraea* | CT | T | intergenic | NA | 2 | 31806962 | 1145225 |
| Sc0000203_707735 | Pool-seq | T+V |  | *Q. petraea* | AT | A | intronic | Qrob_G0237050.2 | 1 | 51985124 | 17848 |
| Sc0000274_909817 | Pool-seq | T+V |  | *Q. petraea* | AG | A | exonic | Qrob_G0320010.2 | 2 | 35418361 | 563863 |
| Sc0000464_236576 | Pool-seq | T | PD | *Q. petraea* | CT | T | exonic | Qrob_G0523500.2 | 5 | 23753953 | 744474 |
| Sc0000481_343721 | Pool-seq | T+V |  | *Q. petraea* | AG | G | intronic | Qrob_G0512850.2 | 5 | 3222156 | 22554 |
| Sc0000974_98303 | Pool-seq | T | PD | *Q. petraea* | AG | A | exonic | Qrob_G0759540.2 | 3 | 47813240 | NA |
| Sc0000485_93093 | Pool-seq | T | VA | *Q. pubescens* | AG | A | exonic | Qrob_G0539160.2 | 1 | 16067626 | 3103379 |
| Sc0000314_149731 | Pool-seq | T+V |  | *Q. pubescens* | AT | A | intergenic | NA | 2 | 55831586 | 5480810 |
| Sc0000314_149731 | Pool-seq | T | VA | *Q. pubescens* | CT | T | intergenic | NA | 7 | 39945935 | 897324 |
| Sc0000047_2398879 | Pool-seq | T+V |  | *Q. pubescens* | AG | A | intronic | Qrob_G0585850.2 | 9 | 16825011 | NA |
| Sc0000062_118505 | Pool-seq | T | DisC | *Q. pubescens* | CT | T | intronic | Qrob_G0091810.2 | 6 | 15935077 | 4083535 |
| Sc0000088_1796044 | Pool-seq | T+V |  | *Q. pubescens* | AC | A | intronic | Qrob_G0328570.2 | 12 | 20880546 | NA |
| Sc0000109_800763 | Pool-seq | T+V |  | *Q. pubescens* | CT | C | exonic | Qrob_G0124910.2 | 1 | 43547799 | 4560446 |
| Sc0000111_693153 | Pool-seq | T+V |  | *Q. pubescens* | CT | T | intergenic | NA | 7 | 28004200 | 286006 |
| Sc0000170_630013 | Pool-seq | T+V |  | *Q. pubescens* | AC | A | intronic | Qrob_G0200770.2 | 6 | 35588545 | 14530676 |
| Sc0000192_329301 | Pool-seq | T+V |  | *Q. pubescens* | AG | A | intronic | Qrob_G0228240.2 | 2 | 39053274 | 2874167 |
| Sc0000482_334917 | Pool-seq | T | PD | *Q. pubescens* | CT | C | intronic | Qrob_G0533790.2 | NA | NA | NA |
| Sc0000403_286465 | Pool-seq | T+V |  | *Q. pyrenaica* | CT | T | intergenic | NA | 2 | 105577540 | 28340724 |
| Sc0000006_2873224 | Pool-seq | T+V |  | *Q. pyrenaica* | AG | A | intronic | Qrob_G0005870.2 | 6 | 21057869 | 5122792 |
| Sc0000014_2037045 | Pool-seq | T+V |  | *Q. pyrenaica* | AC | C | exonic | Qrob_G0064170.2 | 2 | 77236816 | 6145559 |
| Sc0000053_1344456 | Pool-seq | T | DisC | *Q. pyrenaica* | AG | A | intergenic | NA | 6 | 11300343 | 294652 |
| Sc0000085_73024 | Pool-seq | T+V |  | *Q. pyrenaica* | TG | G | intronic | Qrob_G0563240.2 | 10 | 14291154 | 1973542 |
| Sc0000228_1091905 | Pool-seq | T+V |  | *Q. pyrenaica* | TC | C | intergenic | NA | 5 | 35457995 | 7637100 |
| Sc0000269_924931 | Pool-seq | T+V |  | *Q. pyrenaica* | AC | A | intronic | Qrob_G0632320.2 | 2 | 26007366 | 561547 |
| Sc0000287_474090 | Pool-seq | T | AF | *Q. pyrenaica* | AC | C | exonic | Qrob_G0459590.2 | 2 | 71091257 | 6145559 |
| Sc0000307_852597 | Pool-seq | T+V |  | *Q. pyrenaica* | AG | A | intergenic | NA | 7 | 16871124 | 7928330 |
| Sc0000517_383812 | Pool-seq | T+V |  | *Q. pyrenaica* | CG | C | intergenic | NA | 10 | 18200824 | 125219 |
| Sc0000695_157206 | Pool-seq | T | AF | *Q. pyrenaica* | AT | A | exonic | Qrob_G0671270.2 | 8 | 55531952 | 68141 |
| Sc0000778_61930 | Pool-seq | T+V |  | *Q. pyrenaica* | CT | T | intronic | Qrob_G0070130.2 | 10 | 12317612 | 1973542 |
| Sc0000013_2578823 | Pool-seq | T+V |  | *Q. robur* | AG | A | intronic | Qrob_G0010260.2 | 2 | 61312396 | 5480810 |
| Sc0000038_794573 | Pool-seq | T | PQ | *Q. robur* | TG | G | intronic | Qrob_G0701760.2 | 1 | 38987353 | 4560446 |
| Sc0000053_1639108 | Pool-seq | T+V |  | *Q. robur* | AG | A | intergenic | NA | 6 | 11594995 | 256547 |
| Sc0000053_1895655 | Seq-cap | T | DisC | *Q. robur* | CT | T | exonic | Qrob_G0631440.2 | 6 | 11851542 | 256547 |
| Sc0000099_1839376 | Pool-seq | T+V |  | *Q. robur* | AC | A | intronic | Qrob_G0084290.2 | 11 | 6742403 | 2258901 |
| Sc0000111_979159 | Pool-seq | T | AF | *Q. robur* | CT | C | intronic | Qrob_G0135420.2 | 7 | 27718194 | 286006 |
| Sc0000158_462639 | Pool-seq | T+V |  | *Q. robur* | AT | T | exonic | Qrob_G0304430.2 | 2 | 26568913 | 230471 |
| Sc0000158_693110 | Seq-cap | T | AF | *Q. robur* | CG | G | exonic | Qrob_G0304580.2 | 2 | 26799384 | 230471 |
| Sc0000203_689887 | Pool-seq | T+V |  | *Q. robur* | TG | T | intronic | Qrob_G0237030.2 | 1 | 51967276 | 17848 |
| Sc0000225_507799 | Seq-cap | T | AF | *Q. robur* | CG | G | exonic | Qrob_G0487320.2 | 5 | 24498427 | 744474 |
| Sc0000240_289656 | Seq-cap | T | VA | *Q. robur* | AG | G | exonic | Qrob_G0318610.2 | 5 | 4277129 | 1032419 |
| Sc0000270_806328 | Pool-seq | T | AF | *Q. robur* | AG | G | exonic | Qrob_G0692980.2 | 7 | 33545097 | 5540897 |
| Sc0000339_4638 | Pool-seq | T+V |  | *Q. robur* | CT | C | intronic | Qrob_G0473660.2 | 1 | 3718979 | 9245268 |
| Sc0000381_206331 | Seq-cap | T+V |  | *Q. robur* | AC | A | intergenic | NA | 7 | 39048611 | 897324 |
| Sc0000447_521057 | Pool-seq | T+V |  | *Q. robur* | AG | G | exonic | Qrob_G0543330.2 | 10 | 22549567 | 4348743 |
| Sc0000481_366275 | Seq-cap | T | PD | *Q. robur* | CT | T | exonic | Qrob_G0512860.2 | 5 | 3244710 | 22554 |
| Sc0000517_258593 | Pool-seq | T+V |  | *Q. robur* | CT | T | intergenic | NA | 10 | 18075605 | 125219 |
| Sc0000546_456229 | Seq-cap | T | PD | *Q. robur* | AG | G | exonic | Qrob_G0761790.2 | 2 | 35982224 | 563863 |
| Sc0000598_295142 | Seq-cap | T | PD | *Q. robur* | CT | C | exonic | Qrob_G0575620.2 | 2 | 32952187 | 1145225 |
| Sc0000695_225347 | Pool-seq | T+V |  | *Q. robur* | AT | T | intronic | Qrob_G0671240.2 | 8 | 55600093 | 68141 |
| Sc0000796_82698 | Pool-seq | T+V |  | *Q. robur* | AT | T | intronic | Qrob_G0759570.2 | NA | NA | NA |
| Sc0000967_33996 | Pool-seq | T+V |  | *Q. robur* | AT | T | exonic | Qrob_G0709860.2 | 2 | 49368178 | 2526743 |

<sup>a</sup> Study samples (T : Training populations ; V : validation population)
<sup>b</sup> Screening criteria from training to validation (PD : Poor diagnosticity; PQ : Poor quality of cluster delimitation ; Disc : Genotype discrepancy between different multiplexes; VA : variable success (numerous missing data) ; AF : Amplification failure after primer redesign
<sup>c</sup> Chr: Chromosome bearing the diagnostic SNP
<sup>d</sup> Position (in bp) on the chromosome
<sup>e</sup> Distance (in bp) with previous diagnostic SNP on the same chromosome
<sup>f</sup> SNP Identification comprise scaffold number (SC#) and position on the scaffold (Plomion *et al*, 2018)

#### 3.3.3. Allele frequency profiles of diagnostic SNPs in *Q. petraea* populations

We examined the geographic distribution of near-diagnostic alleles between populations within a given species. To illustrate the results we selected populations that are representative of the variation observed among all populations. We first selected a few widely distributed populations that exhibited allele frequencies at all SNPs close to the expected diagnosticity (“EP populations”: Tronçais, Lappwald and Bézange), and added all the populations that deviate from the EP frequency profiles, which we called diverging populations (“DP populations”). The DP populations included three extreme southern populations (Pomieri and Aspromonte in Italy, Montejo in Spain) and one population from the northern distribution edge (Killarney). All the remaining *Q. petraea* populations exhibited frequency profiles similar to the selected EP populations, and are not shown in Table 1 and in Figure 4. While the EP populations exhibited almost full fixation in all near-diagnostic SNPs, the DP populations showed substantial polymorphism (i.e. lower diagnosticity) at a few SNPs in Pomieri and Aspromonte (Sc0000043_1651618, Sc0000135_261350, Sc0000274_909817), and moderate polymorphism distributed among more SNPs in Killarney and Montejo.

**Figure 4.**
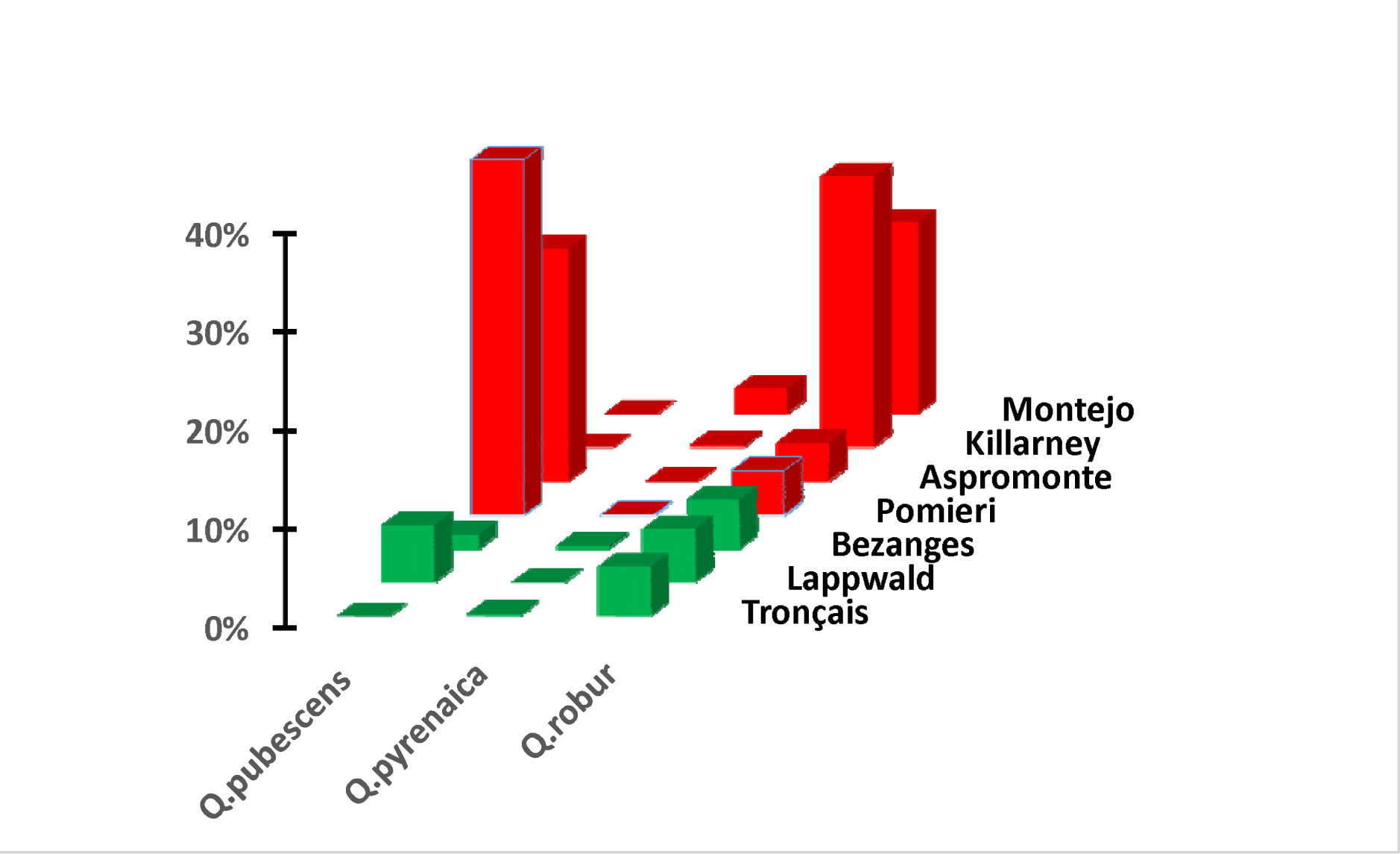
Frequencies of near-diagnostic alleles of *Q.pubescens*, *Q. pyrenaica* and *Q. robur* in *Q*. *petraea* populations. Populations in red and green correspond to DP and EP populations (see text). Shown are the mean frequencies of all near-diagnostic alleles of a given species (*Q.pubescens*, *Q. pyrenaica*, *Q. robur)* in *Q.petraea* populations. Geographic locations of *Q. petraea* populations are shown in Figure 1a.

Additionally, we examined the occurrences of near-diagnostic alleles of the other three species in *Q. petraea* populations (Figure 4). Interestingly the DP *Q. petraea* populations were also diverging in respect to the frequency of near-diagnostic alleles of *Q. pubescens* (Pomieri and Aspromonte), or *Q. robur* (Killarney and Montejo). The EP populations exhibited lower frequencies of near-diagnostic alleles of the other three species (Figure 4).

#### 3.3.4. Allele frequency profiles of near-diagnostic SNPs in *Q. pubescens*, *Q. robur* and *Q. pyrenaica* populations

To illustrate the intraspecific differentiation of near-diagnostic SNPs in the other three species, we followed the same procedure as for *Q. petraea.* We selected for each species two sets of populations: a subset of populations exemplifying the pattern close to full fixation of near-diagnostic loci at all SNPs (EP populations), and the set of diverging populations (DP populations) that exhibited deviations to this trend.

In the case of *Q. pubescens,* the DP populations (Switzerland and Ventoux) were located at the central northern edge of distribution. These deviations were not evenly distributed across the 7 near-diagnostic SNPs of *Q. pubescens*, but restricted to the same loci in the two populations (Table 2). The two populations Switzerland and Ventoux exhibited also higher frequencies of *Q. petraea* near*-*diagnostic alleles, in comparison to the two EP populations (Figure 7 in Appendix).

In the case of *Q. robur*, there were also two DP populations located at the south western (Pedro) and north western margin of the distribution (Roudsea) (Table 3). These two populations comprised also larger frequencies of near-diagnostic alleles of other white oak species (*Q. pubescens* and *Q. pyrenaica* in the case of Pedro; *Q. petraea* in the case of Roudsea) (Figure 8 in Appendix). Finally, in *Q. pyrenaica*, all populations behave as EP populations (data not shown), eg all *Q. pyrenaica* populations exhibited frequency profiles similar to those shown for *Q. pyrenaica* in Figure 3 and Table 7 in Appendix.

**Table 7 in Appendix.** Overall frequencies of near-diagnostic alleles in the validation populations.

| SNP ID | Reference<br>diagnostic<br>species | Near-diagnostic<br>allele | *Q.petraea* | *Q.pubescens* | *Q.pyrenaica* | *Q.robur* | Diagnosticity |
| --- | --- | --- | --- | --- | --- | --- | --- |
| Sc0000254_6223 | *Q. petraea* | T | **0.863** | 0.116 | 0.074 | 0.023 | 0.792 |
| Sc0000121_355205 | *Q. petraea* | C | **0.966** | 0.151 | 0.109 | 0.068 | 0.857 |
| Sc0000043_1651618 | *Q. petraea* | G | **0.738** | 0.156 | 0.023 | 0.027 | 0.669 |
| Sc0000083_147504 | *Q. petraea* | A | **0.876** | 0.121 | 0.320 | 0.046 | 0.714 |
| Sc0000118_1466708 | *Q. petraea* | G | **0.873** | 0.117 | 0.046 | 0.077 | 0.793 |
| Sc0000135_261350 | *Q. petraea* | C | **0.823** | 0.132 | 0.056 | 0.082 | 0.733 |
| Sc0000145_700044 | *Q. petraea* | T | **0.949** | 0.074 | 0.015 | 0.035 | 0.908 |
| Sc0000203_707735 | *Q. petraea* | A | **0.903** | 0.060 | 0.042 | 0.029 | 0.859 |
| Sc0000274_909817 | *Q. petraea* | A | **0.786** | 0.086 | 0.123 | 0.018 | 0.710 |
| Sc0000481_343721 | *Q. petraea* | G | **0.926** | 0.242 | 0.085 | 0.049 | 0.801 |
| Sc0000314_149731 | *Q. pubescens* | A | 0.116 | **0.660** | 0.050 | 0.054 | 0.587 |
| Sc0000047_2398879 | *Q. pubescens* | A | 0.043 | **0.896** | 0.045 | 0.019 | 0.860 |
| Sc0000088_1796044 | *Q. pubescens* | A | 0.027 | **0.906** | 0.000 | 0.007 | 0.895 |
| Sc0000109_800763 | *Q. pubescens* | C | 0.121 | **0.752** | 0.015 | 0.007 | 0.704 |
| Sc0000111_693153 | *Q. pubescens* | T | 0.337 | **0.996** | 0.061 | 0.012 | 0.859 |
| Sc0000170_630013 | *Q. pubescens* | A | 0.015 | **0.657** | 0.000 | 0.005 | 0.650 |
| Sc0000192_329301 | *Q. pubescens* | A | 0.010 | **0.697** | 0.054 | 0.006 | 0.674 |
| Sc0000403_286465 | *Q. pyrenaica* | T | 0.008 | 0.014 | **0.900** | 0.011 | 0.889 |
| Sc0000006_2873224 | *Q. pyrenaica* | A | 0.012 | 0.016 | **0.962** | 0.009 | 0.950 |
| Sc0000014_2037045 | *Q. pyrenaica* | C | 0.006 | 0.000 | **0.908** | 0.004 | 0.905 |
| Sc0000085_73024 | *Q. pyrenaica* | G | 0.004 | 0.024 | **0.946** | 0.000 | 0.937 |
| Sc0000228_1091905 | *Q. pyrenaica* | C | 0.003 | 0.004 | **0.844** | 0.002 | 0.841 |
| Sc0000269_924931 | *Q. pyrenaica* | A | 0.003 | 0.000 | **0.900** | 0.004 | 0.898 |
| Sc0000307_852597 | *Q. pyrenaica* | A | 0.013 | 0.000 | **0.546** | 0.022 | 0.534 |
| Sc0000517_383812 | *Q. pyrenaica* | C | 0.001 | 0.022 | **0.975** | 0.004 | 0.966 |
| Sc0000778_61930 | *Q. pyrenaica* | T | 0.004 | 0.000 | **0.969** | 0.000 | 0.968 |
| Sc0000013_2578823 | *Q. robur* | A | 0.078 | 0.028 | 0.047 | **0.814** | 0.763 |
| Sc0000053_1639108 | *Q. robur* | A | 0.094 | 0.015 | 0.071 | **0.925** | 0.865 |
| Sc0000099_1839376 | *Q. robur* | A | 0.017 | 0.027 | 0.000 | **0.881** | 0.866 |
| Sc0000158_462639 | *Q. robur* | T | 0.043 | 0.000 | 0.054 | **0.799** | 0.767 |
| Sc0000203_689887 | *Q. robur* | T | 0.072 | 0.029 | 0.078 | **0.796** | 0.736 |
| Sc0000339_4638 | *Q. robur* | C | 0.076 | 0.000 | 0.062 | **0.922** | 0.876 |
| Sc0000381_206331 | *Q. robur* | A | 0.054 | 0.000 | 0.108 | **0.941** | 0.887 |
| Sc0000447_521057 | *Q. robur* | G | 0.213 | 0.028 | 0.000 | **0.956** | 0.876 |
| Sc0000517_258593 | *Q. robur* | T | 0.104 | 0.091 | 0.069 | **0.960** | 0.872 |
| Sc0000695_225347 | *Q. robur* | T | 0.140 | 0.008 | 0.008 | **0.921** | 0.869 |
| Sc0000796_82698 | *Q. robur* | T | 0.083 | 0.074 | 0.133 | **0.901** | 0.804 |
| Sc0000967_33996 | *Q. robur* | T | 0.077 | 0.004 | 0.085 | **0.965** | 0.910 |

Frequencies in bold characters correspond to near-diagnostic alleles of the reference diagnostic species

### 3.4. Multilocus structure of near-diagnostic SNPs

We used a principal component analysis (PCA) in the validation populations to assess and illustrate species differentiation (Figure 5). We added 13 samples of known first generation hybrid origin to the species samples. Ten samples resulted from controlled interspecific crosses, and three came from parentage analysis conducted in a mixed *Q. petraea-Q. robur* stand (Truffaut *et al*, 2017). A combination of the three first components allowed to visually differentiate the four different species. While principal component 1 differentiated mainly *Q*. *petraea* and *Q. robur* (Figure 5a), component 3 distinguished *Q. pyrenaica* from the three other species (Figure 5b), and the biplot of component 2 and 3 provided the best visual separation between *Q. pubescens* and *Q. petraea* (Figure 5c).

**Figure 5.**
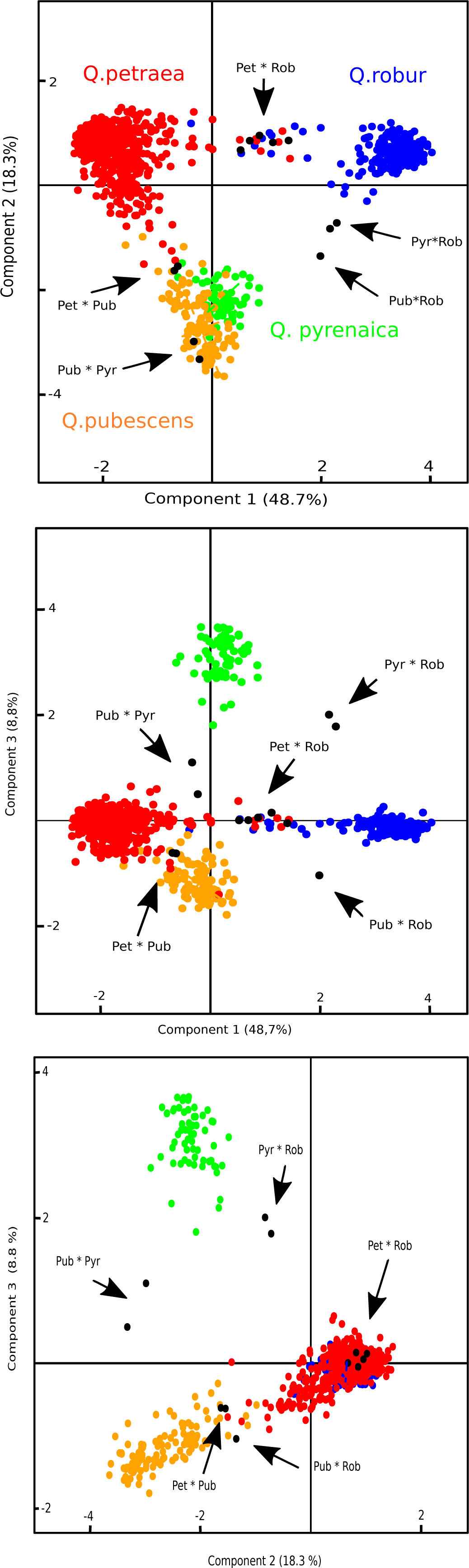
Biplot of principal components of tree samples based on a Principal Component Analysis (PCA) conducted in the validation populations. Figure 1a: Biplot of component 1 and 2 Figure 5b: Biplot of component 1 and 3 Figure 5c: Biplot of component 2 and 3 Numbers between brackets stand for the percentage of variation explained by the component. Red dots: *Q. petraea* samples; Orange dots: *Q. pubescens* samples; Green dots: *Q. pyrenaica* samples; Blue dots: *Q. robur* samples; Black dots: hybrids, Pet*Pub: *Q.petraea*Q.pubescens* hybrids; Pet*Rob: *Q.petraea*Q.robur* hybrids; Pub*Pyr: *Q.pubescens*Q.pyrenaica* hybrids; Pub*Rob: *Q.pubescens*Q.robur* hybrids.

**Figure 6 in Appendix.**
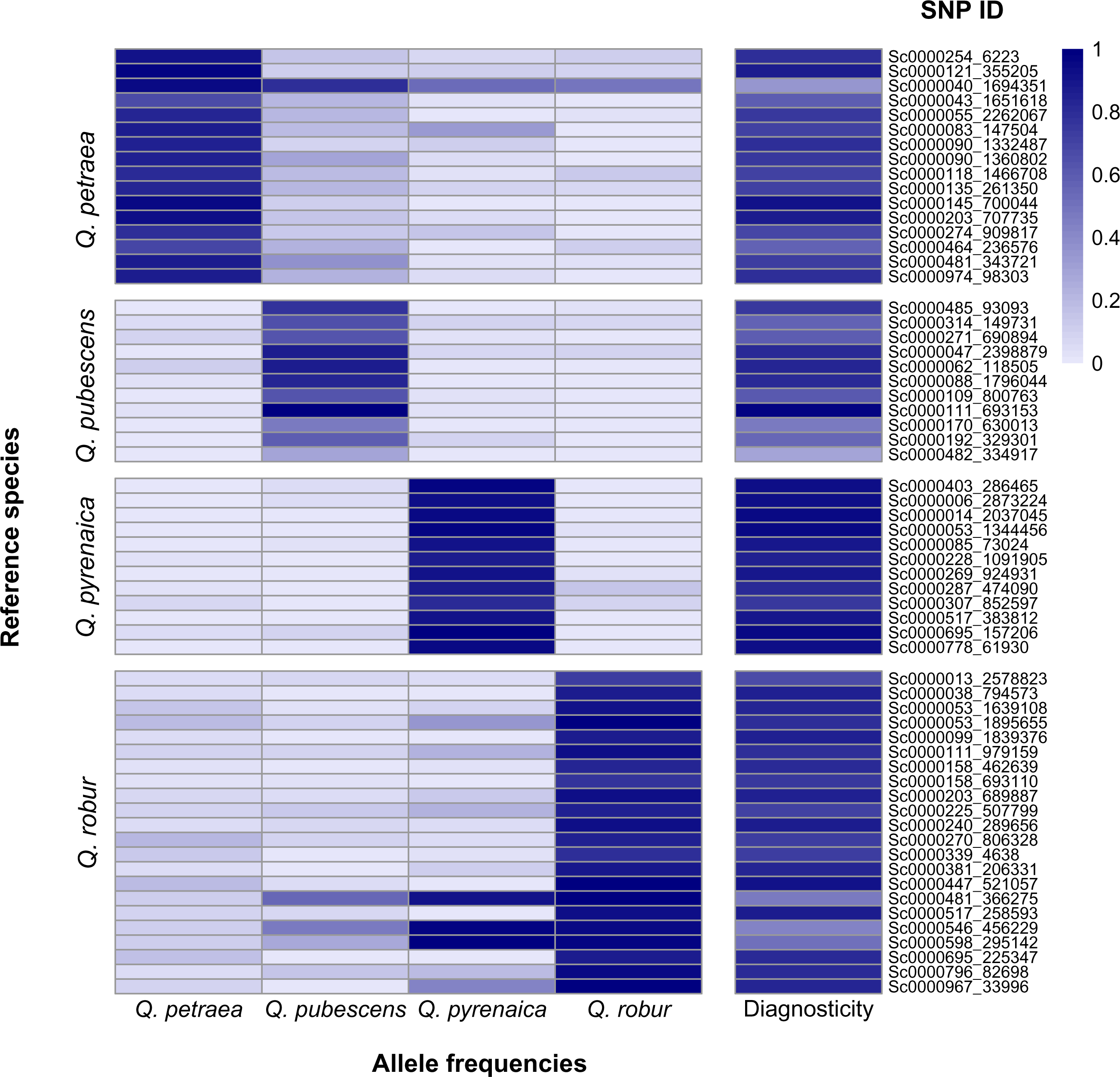
Heat map of frequencies of near-diagnostic alleles and diagnosticity in the training populations of the four species. SNPs are clustered for their diagnostic value for each species (reference species): First to fourth columns correspond respectively to near-diagnostic alleles of *Q*. *petraea*, *Q.pubescens*, *Q. pyrenaica* and *Q. robur*.

These multilocus representations showed that there is a small number of samples located at intermediate positions, especially between *Q. petraea* and *Q. robur* (Figure 5a), and between *Q. petraea* and *Q. pubescens* (Figure 5c). These regions of the PCA are also occupied by known interspecific hybrids, suggesting that the species samples, although identified as pure species in the field, represent either hybrids or introgressed forms. These intermediate positions are also preferentially occupied by trees belonging to diverging populations, as shown by the targeted PCA analysis on the two pairs of species sharing intermediate samples: *Q. petraea* and *Q. pubescens* (Figure 9 in Appendix), *Q. petraea* and *Q. robur* (Figure 10 in Appendix).

## 4. Discussion

We explored large scale existing genomic resources in four European white oaks of the subsection Roburoid (*Q. petraea, Q. pubescens, Q. pyrenaica, Q. robur*) to sceen their genomes for near-diagnostic SNPs that could be used for molecular fingerprinting (species and hybrid identification) in forest research and operational forestry, as wood or seed traceability in the wood chain and in forest nurseries. Despite the widely reported low interspecific genetic differentiation among European white oak species, we were able to identify a subset of SNPs that exhibited near-diagnostic features across their species’ distribution ranges. Moreover, mutlivariate analysis showed that these markers can be used for reliable hybrid detection and accurate quantification of admixture levels. However, diagnosticity varied substantially among species, among populations within species, and among SNPs. In the following, we discuss these variations in relation to the known evolutionary history and genetic interactions among and within the four species.

### 4.1 Variation of diagnosticity among species

Diagnosticity was highest in *Q. pyrenaica* (0.876) and lowest in *Q. pubescens* (0.747) with *Q. robur* and *Q. petraea* showing intermediate values. Near-diagnostic SNPs are likely located in genomic regions that exhibit larger divergence and/or regions prevented from interspecific gene flow. The range of diagnosticity among the four species, may therefore reflect the variation of divergence time and/or the variation of the intensity of gene flow during the ongoing interglacial period.

It is striking to notice that higher and lower diagnosticity was observed for species that showed the older (*Q. pyrenaica, Q. robur*) and more recent (*Q. petraea, Q. pubescens*) divergence, respectively (Leroy *et al*, 2017). Fixation of near-diagnostic SNPs in species with large population sizes as in oaks requires long time periods. Consequently, lower diagnosticity is likely associated with species that diverged more recently. This is illustrated by *Q. pubescens*, which shows lower diagnosticity due to the higher sharing of near-diagnostic alleles with *Q. petraea* than with the other two species (Figure 3 and Figure 6 in Appendix). Diagnosticty may in addition be dependent on the variation of population size (*Ne*) among species and along divergence, for which we lack any estimation today. Our results may therefore be revisited in the light of future evidence of *Ne* differences. Regarding gene flow, we showed earlier that the four species came into contact only recently, during the late last glacial maximum, after being isolated for most of their earlier history (Leroy *et al*, 2020b; Leroy *et al*, 2017), resulting in gene flow among species. While interfertility among the four species has been shown experimentally by controlled crosses (Lepais *et al*, 2013), hybridization *in natura* has also been observed among the four species in rare mixed forests where all four species co-occur (Lepais and Gerber, 2011; Lepais *et al*, 2009). Interspecific matings of *Q. pyrenaica* in controlled crosses with the remaining three species were quite successful, however occurrences of natural hybridization were less frequent due to the very late flowering of *Q. pyrenaica* in comparison to the three other species (Lepais and Gerber, 2011; Lepais *et al*, 2013). Furthermore *Q. pyrenaica* is mainly distributed in south western Europe, where the other three species are only present in scattered forests, leading, for example, to reported but rare hybridization with *Q. petraea* (Valbuena-Carabana *et al*, 2005) and *Q. robur* (Moracho *et al*, 2016). Altogether, phenological prezygotic barriers and limited overlapping distributions with the other three species may have contributed to reduced genetic exchanges between *Q. pyrenaica* and the other three species, and thus account for the high diagnosticity of the SNPs in of *Q. pyrenaica.* In contrast to *Q. pyrenaica,* no reproductive barriers were observed in *Q. pubescens* when crosses were made with *Q. petraea* as female parent, as interspecific crosses were as successful as intraspecific crosses (Lepais *et al*, 2013). Reduced barriers between these two species were corroborated by frequent admixture detected in genetic surveys conducted in mixed stands of *Q. pubescens* and *Q. petraea* (Alberto *et al*, 2010; Neophytou, 2014; Reutimann *et al*, 2023). As a result, near-diagnostic SNPs of *Q. pubescens* and *Q. petraea* were more frequently shared between the two species (Figure 3 and Figure 6 in Appendix) thus contributing to reduced diagnosticity. Finally interspecific gene exchanges involving *Q. robur* were mainly investigated with regard to *Q*. *petraea.* Uneven gene flow has been repeatedly observed in mixed stands with limited pollination from *Q. robur* to *Q. petraea* (Bacilieri *et al*, 1996; Lagache *et al*, 2013; Lepais *et al*, 2013), with a few exceptions in stands of unbalanced mixtures (Gerber *et al*, 2014). Uneven and unidirectional gene exchanges between these two species may have resulted in higher diagnosticity of *Q. robur* in comparison to *Q. petraea*.

### 4.2 Variation of diagnosticity among populations

There are striking differences of species diagnosticity of the markers among populations within species (Table 1, 2 and 3). In populations of *Q. petraea, Q. pubescens* and *Q. robur* located in the central part of their distributions, high levels of diagnosticity (mean values of SNP diagnosticity of the population) could be observed, while in populations located at the margins of the distributions, southern as well as northern, lower diagnosticity was found. We further showed that populations located at the edges of distribution are characterized by higher frequencies of near-diagnostic alleles of the other three congeneric species, suggesting extensive genetic exchanges (Figure 4, Figure 7 and 8 in Appendix). More frequent interspecific gene flow at the northern edge of distribution has been shown earlier in the case *Q. petraea* and *Q. robur* (Beatty *et al*, 2016; Jensen *et al*, 2009; Gerber *et al*, 2014), and has been interpreted as a driver of the succession dynamics at the northern colonization front of the two species (Kremer and Hipp, 2020; Petit *et al*, 2003). In our study, the sessile oak population Killarney (Figure 4) and the pedunculate oak population Roudsea (Figure 8 in Appendix) are typical examples illustrating interspecific gene flow between the two species. Similar observations of more frequent hybridization were made in the case of *Q. petraea* and *Q. pubescens* at the northern edge of distribution of *Q. pubescens* (Neophytou *et al*, 2015; Reutimann *et al*, 2020), which may have as well contributed to the expansion of *Q. pubescens*.

**Figure 7 in Appendix.**
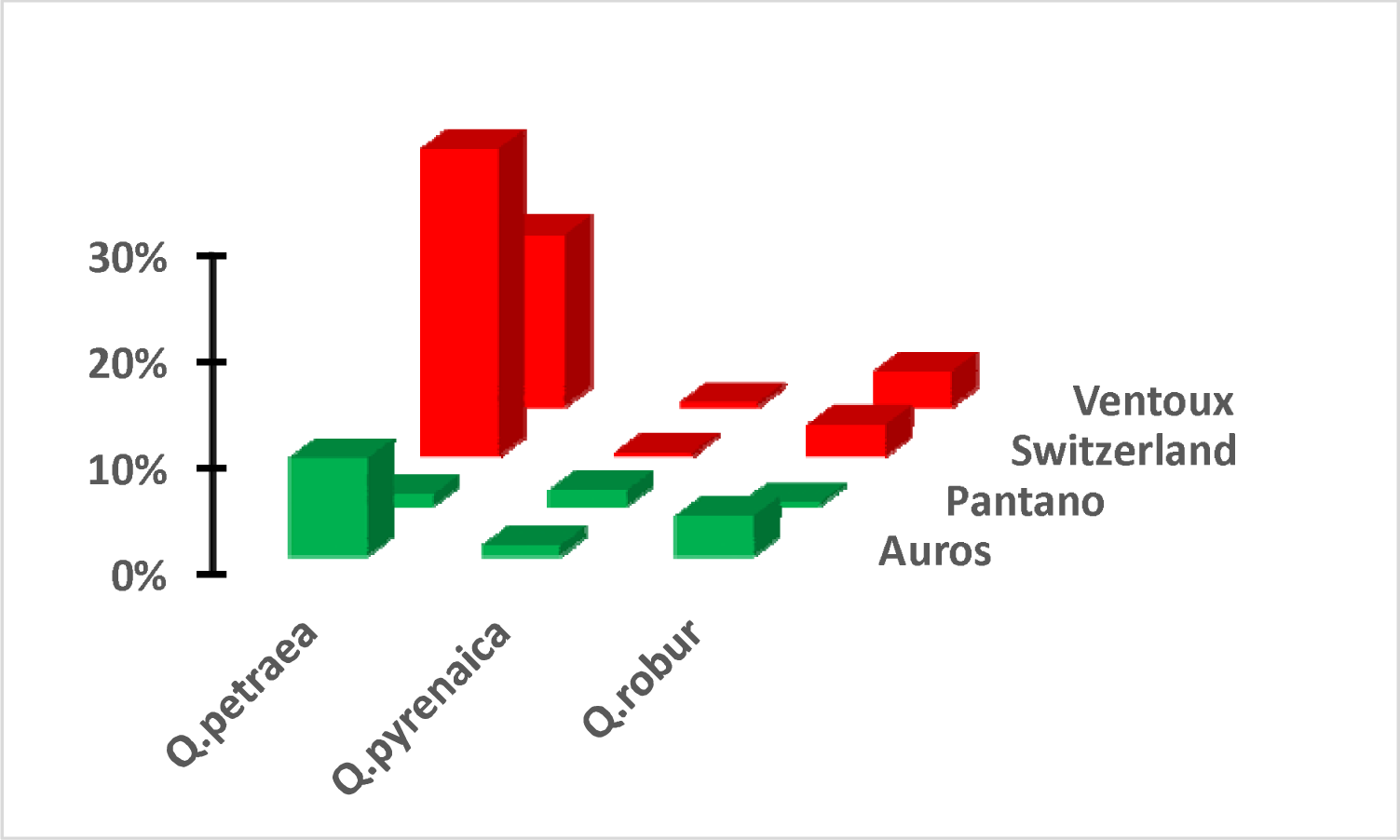
Frequencies of near-diagnostic alleles of *Q.petraea*, *Q. pyrenaica* and *Q. robur* in *Q*. *pubescens* populations. Populations in red and green correspond to DP (diverging populations) and EP (expected populations, see text). Shown are the mean frequencies of all diagnostic alleles of a given species (*Q. petraea, Q. pyrenaica, Q. robur*) in *Q.pubescens* populations. Geographic locations of *Q. pubescens* populations are shown in Figure 1b

**Figure 8.**
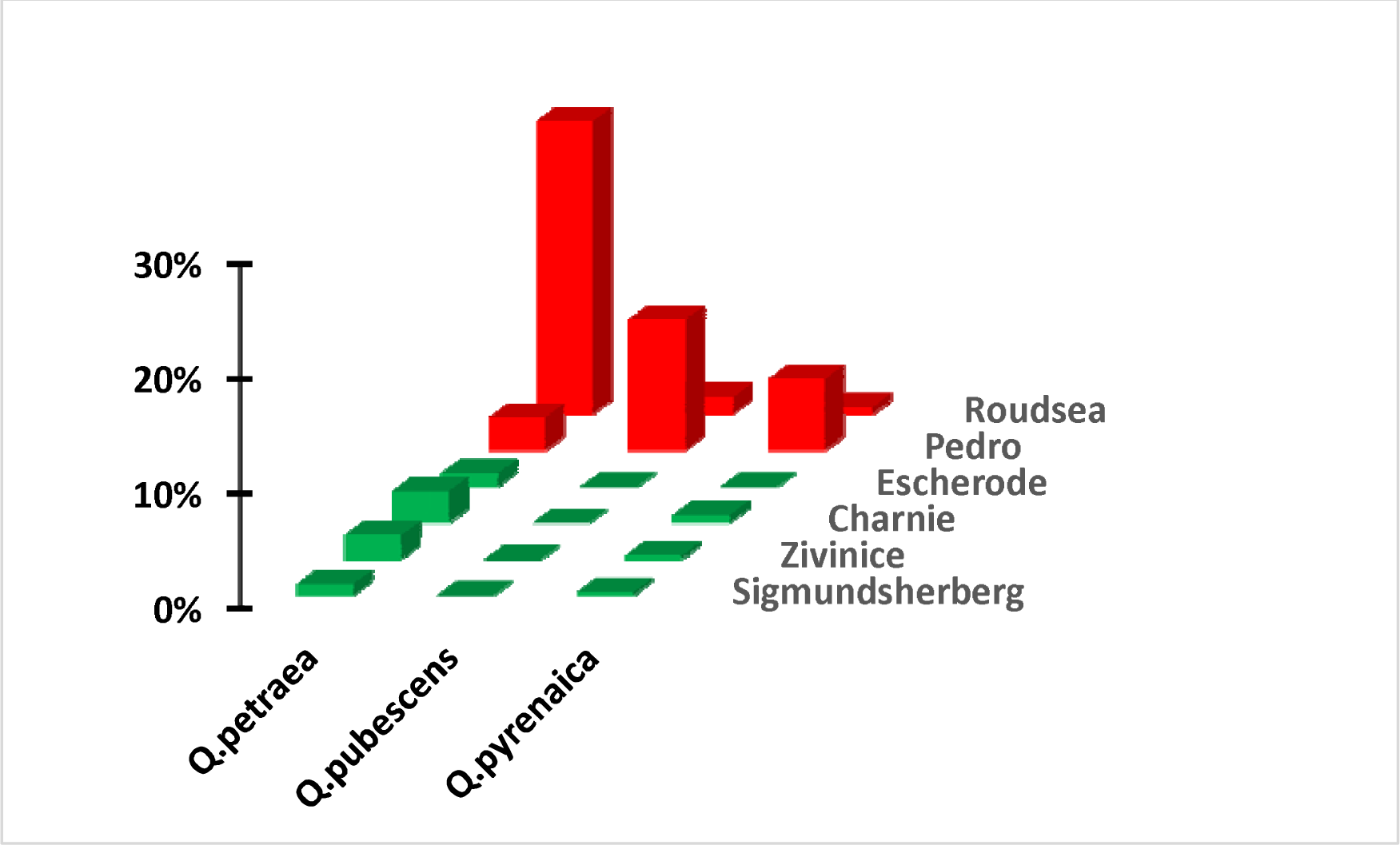
in Appendix Frequencies of near-diagnostic alleles of *Q.petraea*, *Q. pubescens*, and *Q. pyrenaica* in *Q*. *robur* populations. Populations in red and green correspond to DP (diverging populations) and EP (expected populations, see text). Shown are the mean frequencies of all diagnostic alleles of a given (*Q.petraea, Q. pubescens, Q.pyrenaica*) species in *Q.robur* populations. Geographic locations of *Q.robur* populations are shown in Figure 1d

**Figure 9 in Appendix.**
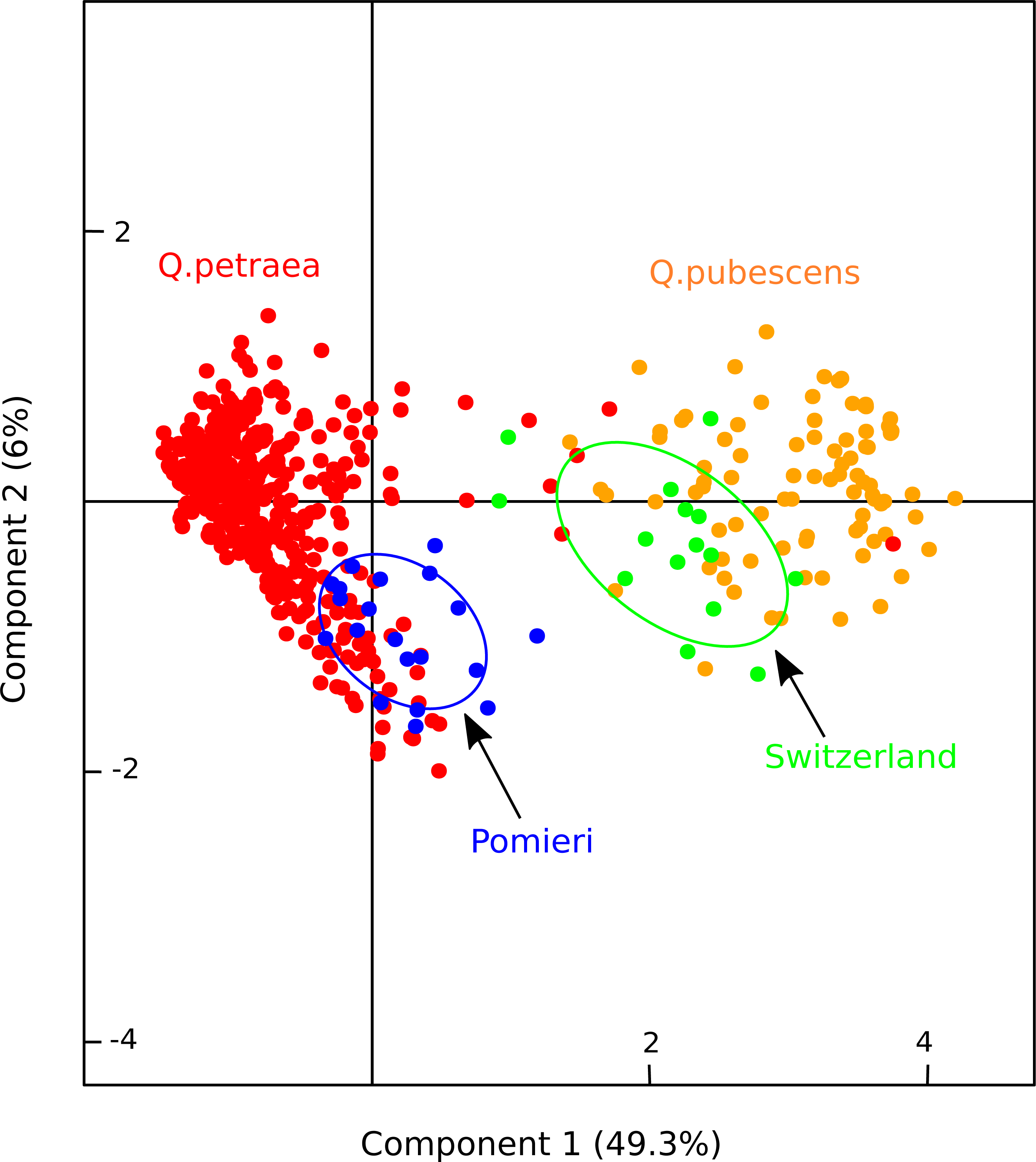
Biplot of principal components of tree samples based on a Principal Component Analysis (PCA) conducted in the *Q. pubescens* and *Q. petraea* validation populations. Red dots: *Q. petraea* samples; Orange dots: *Q. pubescens* samples. Blue dots: *Q. petraea* Pomieri population. Green dots: *Q. pubescens* Switzerland population

**Figure 10 in Appendix.**
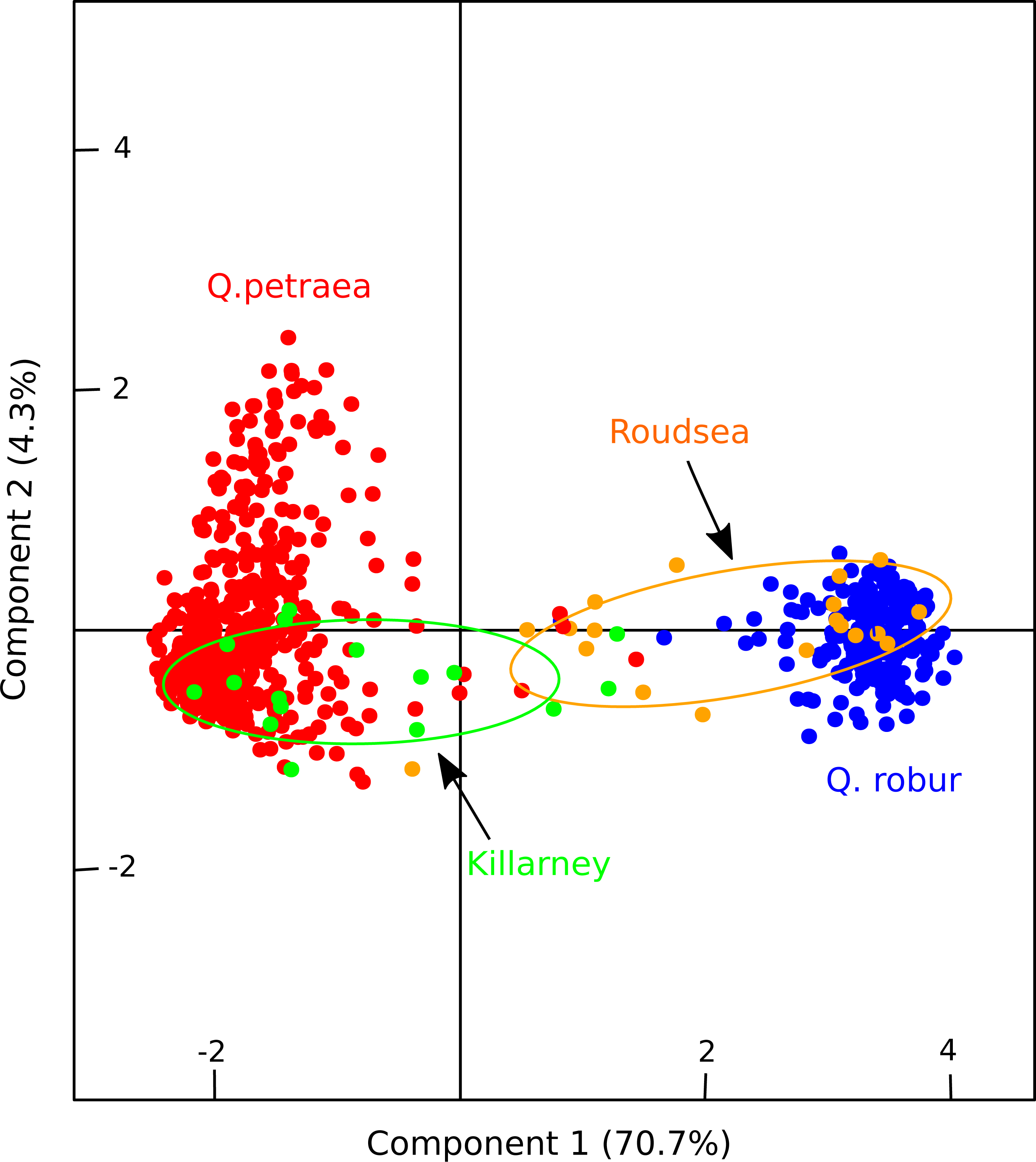
Biplot of principal components of tree samples based on a Principal Component Analysis (PCA) conducted in the *Q. robur* and *Q. petraea* validation populations. Red dots: *Q. petraea* samples; Blue dots: *Q. robur* samples. Green dots: *Q. petraea* Killarney population. Orange dots: *Q.robur* Roudsea population

In populations located at the southern edge of distribution (Pomieri, Aspromonte, and Montejo for *Q. petraea*, Figure 1 and Figure 4), the lower diagnosticity may have resulted from more ancient genetic exchanges with *Q. pubescens* and *Q. robur.* Indeed the two italian populations (Pomieri and Aspromonte) in Sicilia and Calabria consist today in almost pure stands, where *Q. pubescens* is extremely rare, if not absent (Bagnato *et al*, 2012; Modica, 2001), while our results indicated introgression of *Q. pubescens* into *Q. petraea* (Figure 4). Similarly the sessile oak population Montejo, in central Spain, is introgressed by *Q. robur* (Figure 4), where the latter species is absent today and where contemporary hybridization has rather been detected with *Q. pyrenaica* (Valbuena-Carabana *et al*, 2005). Finally, a similar scenario holds for the pedunculate oak population Pedro, which is located at the extreme southern edge of distribution of *Q. robur* (Figure 1; Table3, Figure 8 in Appendix). Hybridization has been observed with *Q. pyrenaica* which is today the most frequent species in the area (Moracho *et al*, 2016) and is confirmed by our results revealing the presence of *Q. pyrenaica* near-diagnostic alleles in the *Q. robur* population (Figure 8 in Appendix). However, introgression by *Q. pubescens* is even more pronounced in our data despite the today’s absence of *Q. pubescens* in Extremadura (Figure 8 in Appendix). To sum up, when comparing our results with previous investigations on interspecific gene flow, recent and/or ancient gene exchanges have faded diagnosticity in the so-called diverging populations, which are located at the northern or southern margins of the distribution.

### 4.3 Variation of diagnosticity among SNPs

Frequency profiles of near-diagnostic alleles differed markedly across SNP in diverging populations. There were cases where lack of diagnosticity affected mainly the same limited number of loci in a given species (Aspromonte and Pomieri in *Q. petraea*, Table 1; Pedro in *Q. robur*, Table 3; and to a smaller extend Switzerland and Ventoux in *Q. pubescens*, Table 2). In the remaining diverging populations (Killarney for *Q. petraea*, Table 1; and Roudsea for *Q. robur*, Table 3), reduced diagnosticity is more evenly distributed across more if not all loci. Contrasting diagnosticity distribution across loci may likely correlate to the timing of hybridization and introgression among the congeneric species. Recent gene exchanges, as first generation hybridization and subsequent backcrosses will indistinctly impact all loci during the early phase of secondary contact among species, and result in reduced diagnosticity of alleles in sympatric species. Such a scenario may hold for the two northern *Q. petraea* (Killarney) and *Q. robur* (Roudsea) populations. Continuous gene exchanges over multiple generations may ultimately result in heterogeneous genomic landscapes, shaped by variable permeability to gene flow along the chromosomes due to the presence of prezygotic or postzygotic barriers and the heterogeneous recombination landscape. This scenario leads ultimately to the maintenance of near-diagnostic loci in genomic regions impermeable to gene flow, while the remaining part of the genome will become poorly differentiated. While this scenario was supported by ABC simulations (Leroy *et al*, 2020b; Leroy *et al*, 2017), our results further suggest that the genomic distribution of near-diagnostic loci is environment-dependent. It is striking that a very limited number of near-diagnostic alleles discovered in western populations of *Q. petraea* show poor diagnosticity in the southern populations Pomieri and Aspromonte (Table 1). Our results further indicated that this low diagnosticity may be due to more interspecific gene flow with *Q. pubescens,* which suggest preferentially introgression in specific genomic regions–whether adaptive or not-resulting ultimately in heterogeneous genomic distribution of near-diagnostic SNPs especially in marginal range parts. In a recent paper we showed that introgressed regions between *Q. robur* and *Q. petraea* may be more frequent at higher altitudes (Leroy *et al*, 2020a) while in another case study in two Asian oak species the authors found that the genomic landscape of introgression changed in different ecological settings (Fu *et al*, 2022). A similar picture holds for the diverging southern *Q. robur* population Pedro, where diagnosticity is substantially reduced at a few near-diagnostic SNPs in comparison to other *Q. robur* populations (Table 3), most likely due to introgression by *Q. pubescens* and *Q. pyrenaica* (Figure 8 in Appendix). Anecdotally the diverging status of Aspromonte, Pomieri and Pedro echoes with the taxonomic subspecies status that has been assigned to the Sicilian and Calabrian *Q. petraea* populations (*Q. petraea* ssp *austrothyrrenica*, Bagnato *et al*, 2012; Lupini *et al*, 2019; Merlino *et al*, 2014) and to the extreme southern spanish *Q. robur* populations (*Q. robur* ssp *estremadurensis,* Vazquez-Pardo *et al*, 2009).

## Conclusions and outlook

Here we showed that near-diagnostic marker development for species identification is feasible despite few species barriers, extensive secondary contact, and, consequently, frequent hybridization and introgression. Recently we demonstrated that the set of near-diagnostic markers resolved species assignment on fossil and archeological oak wood remains, where anatomical features do not allow to discriminate the four deciduous species (Wagner *et al*, 2023). With the steadily ongoing availability of whole genomes in non model species including oaks (Lazic *et al*, 2021), the search of near-diagnostic markers could be extended to the whole Roburoid subsection facilitating white oak species assignment throughout Europe, beyond the subset of four species that we considered here.The near-diagnostic SNPs for the four white oak species could not only be used in forest research and management for reliable and affordable species assignment, but also to identify admixed individuals and accurately quantify admixture levels in natural populations (Reutimann *et al,* 2020). Because the presented alleles are often almost fixed for the target species, these SNPs also allow the identification of hybrid state (F1, F2, backcrosses, later generation hybrids, etc.) with methods like NEWHYBRIDS (Anderson 2008), and altogether help to understand the importance of hydribization and introgression in evolutionary processes. Together with prospect of emergence of field-based genotpying techniques (Urban *et al*, 2021), such near-diagnostic markers would even allow fast fingerprinting *in-situ* to make decision for forest managers and scientists.

## Declarations

### Ethics approval and consent to participate

not applicable

### Consent for publication

not applicable

### Availability of data

The data that support the findingd of this study will be available at the publicly accessible data repository of INRAE: The url address will be completed after the manuscript is accepted for publication.

### Competing interests

there are no competing interests

## Funding

This research was supported by the European Research Council through an Advanced Grant (project TREEPEACE # FP7-339728), by an ANR (Agence Nationale de la Recherche) Grant (project GENOAK 2022, #BSV6-009-02), and by the EVOLTREE Opportunity call (project OakID2).

## Authors’ contributions

Conception of the study: TL, AK; Sampling and collection of material: AK, CR, TL; Discovery of near-diagnostic markers in pool-sequenced resources: TL; Discovery of near-diagnostic markers in sequence captured resources: IL; Design of multiplexes and genotyping of natural populations: EG, AD; Data analysis: AK, TL, SW; Writing of the manuscript: AK, EG, TL. All authors reviewed the manuscript.

## Acknowledgements

We thank colleagues that contributed to the collection of material made for this study: Dalibor Ballian (Bosnia and Herzegovina), María Valbuena Carabaña and Luis Gil (Spain), Giovanni Giuseppe Vendramin (Italy). We extent our appreciation to partners of the former EU supported FAIROAK and OAKFLOW projects, and of the EVOLTREE Network of Excellence, who collected material included in this study. The MassArray genotyping was performed at the PGTB (doi:10.15454/1.5572396583599417E12) with the help of Laure Dubois, Céline Lalanne and Marie Massot. We are grateful to François Ehrenmann for his contributions to the figures of the manuscript.

## Key message

Mining genome-wide DNA sequences enabled the discovery of near-diagnostic markers for species assignment in European white oaks despite their low interspecific differentiation.

## Notes

### Competing Interest Statement

The authors have declared no competing interest.

